# Ectopic activation of the Spindle Assembly Checkpoint reveals its biochemical design and physiological operation

**DOI:** 10.1101/154054

**Authors:** Chu Chen, Ian P. Whitney, Anand Banerjee, Palak Sekhri, David M. Kern, Adrienne Fontan, John J. Tyson, Iain M. Cheeseman, Ajit P. Joglekar

**Affiliations:** Department of Biophysics, University of Michigan, Ann Arbor, MI 48109; Whitehead Institute for Biomedical Research, and Department of Biology, MIT, Nine Cambridge Center, Cambridge, MA 02142; Department of Biological Sciences, Virginia Polytechnic Institute & State University, Blacksburg, VA 24061; Cell & Developmental Biology, University of Michigan Medical School, Ann Arbor, MI 48109

**Keywords:** cell cycle, mitosis, signaling, mitotic checkpoint, kinetochore

## Abstract

Switch-like activation of the Spindle Assembly Checkpoint (SAC) is critical for accurate chromosome segregation during cell division. To determine the mechanisms that implement it, we engineered an ectopic, kinetochore-independent SAC activator, the “eSAC”. The eSAC stimulates the SAC signaling cascade by artificially dimerizing the Mps1 kinase domain and a cytosolic KNL1 phosphodomain, the signaling scaffold in the kinetochore. Quantitative analyses and mathematical modeling of the eSAC reveal that the recruitment of multiple SAC proteins by the KNL1 phosphodomain stimulates synergistic signaling, which enables a small number of KNL1 molecules produce a disproportionately strong anaphase-inhibitory signal. However, when multiple KNL1 molecules signal concurrently, they compete for a limited cellular pool of SAC proteins. This frustrates synergistic signaling and modulates signal output. Together, these mechanisms institute automatic gain control – inverse, non-linear scaling between the signal output per kinetochore and the unattached kinetochore number, and thus enact the SAC switch.

## Introduction

Accurate chromosome segregation during cell division requires that the sister kinetochores on each replicated chromosome stably attach to spindle microtubules before the cell divides. If one or more kinetochores are unattached, they activate a biochemical signaling cascade known as the Spindle Assembly Checkpoint (SAC) (Foley and Kapoor, 2013). This cascade produces an anaphase-inhibitory signal known as the ‘Mitotic Checkpoint Complex’ (MCC). The MCC inhibits the degradative activity of the Anaphase Promoting Complex/Cyclosome (APC/C) throughout the cell, and thus prevents anaphase onset (Musacchio, 2015).

The SAC behaves like a switch with respect to the presence of unattached kinetochores in the dividing cell: it is ‘ON’, if the cell contains one or more unattached kinetochores, and ‘OFF’ otherwise (Figure 1A). The realization of an on-off response using a biochemical signaling cascade can be challenging, especially in the case of the SAC. The SAC signaling cascade is stimulated by unattached kinetochores, which are complex and still incompletely defined nanoscopic structures. Each unattached kinetochore contains a fixed number of stably bound molecules of the protein KNL1 (~ 250 molecules of KNL1 per human kinetochore, (Suzuki et al., 2015)) that serves as the physical scaffold for SAC signaling. For the switch-like activation of the SAC, a single unattached kinetochore must recruit multiple copies of several different signaling proteins from the cytosol using the small number of KNL1 molecules, and generate a sufficiently large flux of the MCC to inhibit APC/C throughout the entire volume of the cell. If the design of the SAC does not enable the kinetochore to do so, the result is higher rates of chromosome missegregation. The large number of unattached kinetochores in a prophase cell poses an additional challenge to the biochemical design of the SAC. For example, there are ~ 92 unattached, signaling kinetochores in a human cell in prophase, but this number decreases rapidly until only one unattached kinetochore remains (Figure 1A). Ideally, the biochemical design of the SAC should ensure that the large number of unattached kinetochores does not generate an excessively strong anaphase-inhibitory signal, but a single unattached kinetochore does produce the signal necessary to delay anaphase onset (Figure 1A, red curve in bottom graph). If the SAC meets both these characteristics, then the dividing cell can maximize the accuracy of chromosome missegregation and minimize unnecessary delays in cell division. However, how the SAC signaling cascade responds to the number (i.e. the dosage) of unattached kinetochores in the dividing cell, and the mechanisms that shape this response remain unclear.

**Figure 1.**
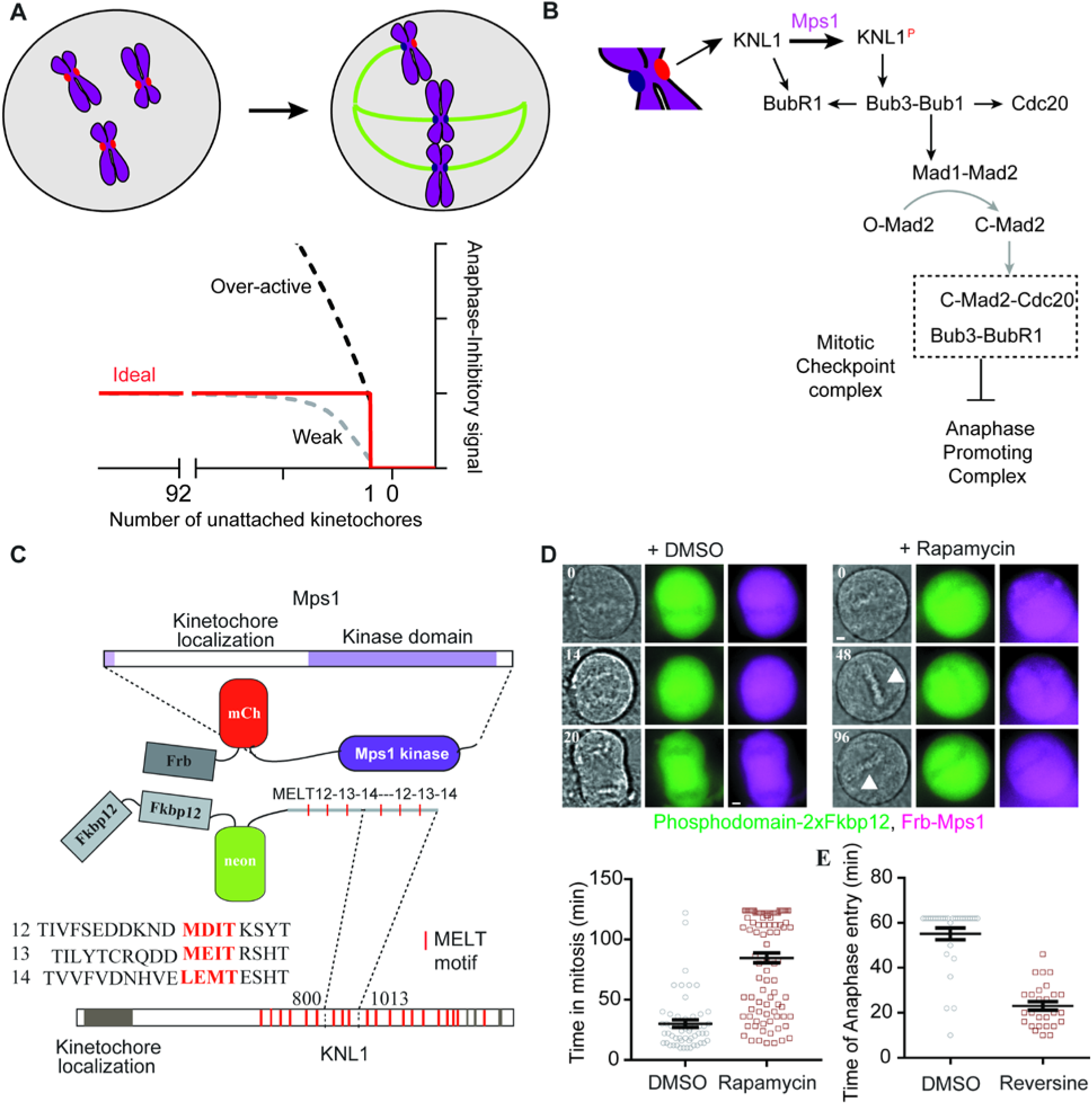
Cytosolic dimerization of the Mps1 kinase domain and a minimal KNL1 phosphodomain is sufficient to induce metaphase arrest. **(A)** Cartoon schematic of the mitotic progression of a cell from prophase (left) to prometaphase (right). Bottom: possible responses of the SAC signaling cascade to the changing number of unattached kinetochores in a human cell. (B) Diagram of the SAC signaling cascade. Black arrows imply protein recruitment to the kinetochore, gray arrows mark cytoplasmic reactions. (**C**) Scheme for conditional dimerization Mps1 with the minimal KNL1 phosphodomain. (**D**) Top, Bright-field and fluorescence images of HeLa cells from time lapse imaging display the indicated proteins. Elapsed time (minutes) indicated in the top left corner. Scale bar ~ 2.4 microns. Bottom, Duration of mitosis in a 2-hour time-lapse experiment (n=55 and 98 for DMSO and Rapamycin respectively). (**E**) Time until anaphase entry after treatment with either DMSO (n=30) or Reversine (n=27, ≥2 independent trials) of cells in rapamycin-induced arrest.

Defining the mechanisms that enable the switch-like operation of the SAC has been challenging, in part because of the complexity of the SAC signaling cascade (Figure 1B). Current models suggest that an unattached kinetochore activates the SAC by allowing the Mps1 kinase to phosphorylate the kinetochore protein KNL1/Spc105 at phosphorylation sites known as the ‘MELT motifs’ due to their consensus sequence (Figure 1A) (Hiruma et al., 2015; Ji et al., 2015; London et al., 2012; Vleugel et al., 2013). Each phosphorylated MELT motif can directly bind to one molecule of the Bub3-Bub1 complex or the Bub3-BubR1 complex (Primorac et al., 2013; Zhang et al., 2016). The Bub1 molecules recruited to kinetochores are also phosphorylated by Mps1, which allows them recruit the Mad1-Mad2 complex (London and Biggins, 2014). The Bub3-BubR1 complex is also recruited via Bub 1-BubR1 heterodimerization (Overlack et al., 2015). BubR1 binds to the APC activator Cdc20 (Di Fiore et al., 2015). The Mad1-Mad2 complex then facilitates the conformational conversion of the inactive ‘Open-Mad2’ into the active ‘Closed-Mad2’ (Musacchio, 2015). In this manner, the unattached kinetochore recruits the MCC subunits, and primes the generation of the MCC, which is the diffusible anaphase-inhibitory signal (Figure 1A-B). These complex, interdependent signaling reactions are difficult to quantify *in vivo*. Moreover, their localization within the nanoscopic kinetochore structure makes a simple analysis based on the mass action law problematic, because the crowded environment of these reactions is likely to influence reaction rates significantly. Finally, the kinetochore-localized signaling activity also makes the definition of the dose-response characteristics for the SAC signaling cascade impractical, because this entails the daunting task of generating specific numbers of unattached kinetochores (Dick and Gerlich, 2013). Owing to these challenges to obtaining quantitative data, the biochemical design and physiological operation of the SAC has remained poorly understood.

To elucidate the biochemical design of the SAC, we engineered a simple method to hijack control of the SAC from the kinetochore. We refer to this ectopic SAC activator as the “eSAC”. The eSAC uses controllable heterodimerization of a cytosolic fragment of the KNL1 phosphodomain containing a specified number of MELT motifs and the Mps1kinase domain. We designed this system to ensure that neither protein is able to localize to kinetochores so that eSAC activity independent of kinetochores. Using the eSAC, we show that the phosphorylation of the MELT motifs by Mps1 is sufficient to recruit SAC proteins to the cytosolic KNL1 phosphodomain, generate the MCC, and delay anaphase onset. Crucially, quantitative analyses of the effect of the eSAC on the duration of mitosis reveal the dose-response characteristics of the SAC signaling cascade with high resolution.

## Results

### Engineering a controllable, ectopic SAC activator or ‘eSAC’

At unattached kinetochores, the phosphorylation of MELT motifs by Mps1 is necessary for SAC signaling. Therefore, we reasoned that we could bypass the involvement of the kinetochore in SAC activation by bringing Mps1 into direct contact with KNL1 in the cytosol in human cells (Aravamudhan et al., 2015). To construct such a kinetochore-independent SAC activator, we first identified a domain of KNL1 that is unable to localize to kinetochores. For this, we ectopically expressed fluorescently labeled fragments of KNL1 in HeLa cells (Figure S1). As expected, the known kinetochore-localization domain at the C-terminus of KNL1 robustly localized to kinetochores (Petrovic et al., 2014). Surprisingly, regions containing the N-terminus of KNL1 also localized to kinetochores transiently in prometaphase. The N-terminus of KNL1 contains the “KI” motifs, which can constitutively bind Bub1 and BubR1 (Kiyomitsu et al., 2011; Krenn et al., 2014; Krenn et al., 2012). However, these motifs are not essential for SAC signaling (Vleugel et al., 2013). Therefore, we selected a region in the center of the KNL1 phosphodomain containing the 12^th^, 13^th^, and 14^th^ MELT motif in KNL1 (a.a. 800-1013, Figure 1C; characterized in (Vleugel et al., 2013)). We fused neonGreen-FKBP12 to the C-terminus of either one or two copies of this region to obtain two versions of the phosphodomain containing either 3 or 6 MELT motifs. Hereafter, we refer to these minimal KNL1 phosphodomains as ‘eSAC phosphodomains’. We fused FRB-mCherry to the N-terminus of full-length Mps1 kinase (Figure 1C). Next, we created stable HeLa cell lines by integrating a bicistronic cassette containing the genes encoding these fusion proteins at an engineered site in the HeLa genome using a previously established method (Figure S1C) (Ballister et al., 2014). In these cell lines, the eSAC phosphodomain is expressed constitutively, whereas the FRB-fused Mps1 kinase is conditionally expressed by a *Tet_ON_* promoter (Figure S1D).

When we induced the dimerization of the eSAC phosphodomain with the Frb-fused Mps1 kinase by adding rapamycin, cells displayed a prolonged mitotic arrest (Figure 1D; Experimental Procedures). These cells maintained an aligned metaphase plate indicating that they were unable to initiate anaphase even though kinetochore-microtubule interactions were intact (Figure 1C, arrowhead). Furthermore, the observed arrest was reversed rapidly upon the inhibition of Mps1 kinase activity by the small molecule inhibitor Reversine demonstrating that the observed arrest depends on Mps1 kinase activity (Figure 1E) (Santaguida et al., 2010). Thus, the observed arrest was indicative of ectopic SAC activation.

In these experiments, we dimerized full-length Mps1 with the eSAC phosphodomain. Therefore, even though the eSAC phosphodomain could not localize to kinetochores, it remained possible that the Frb-fused full-length Mps1 could do so and involve kinetochores in its activity. To ensure completely kinetochore-independent operation of the eSAC, we removed both kinetochore-localization domains of Mps1 (Figure 2A) (Hiruma et al., 2015; Ji et al., 2015). We refer to this minimal Mps1 kinase domain as the eSAC kinase domain. Rapamycin-induced dimerization of the eSAC kinase domain with the eSAC phosphodomain produced a prolonged metaphase arrest with an average duration that comparable to duration of the mitotic arrest induced by treatment with the microtubule stabilizing drug taxol at 100 nM (Figure 2A, also see Supplemental Movies S1-S3). Importantly, neither of the two eSAC proteins could be detected at kinetochores in these metaphase-arrested cells (Figure 2A).

**Figure 2.**
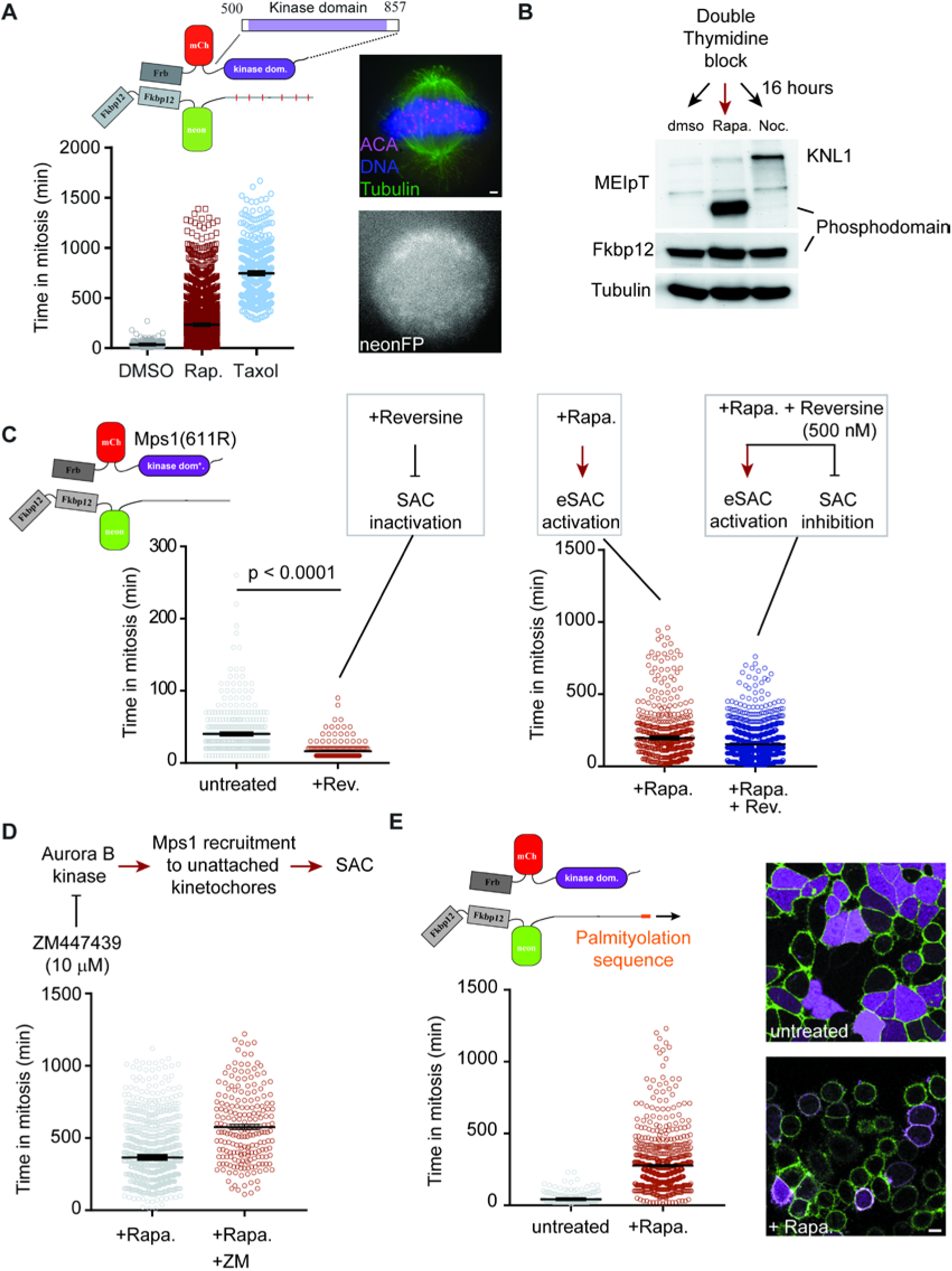
eSAC induces metaphase arrest by acting independently of kinetochores. (**A**) Top, eSAC schematic. Left, Duration of mitosis for untreated and rapamycin-treated cells (n=629 and 2705 respectively). Top right, Psuedocolored fluorescence image of a cell in rapamycin-induced metaphase arrest stained as indicated, Bottom right: absence of kinetochore localization of the phosphodomain visualized by neon fluorescence (scale bar ~ 1.2 μm). Horizontal lines indicate mean ± s.e.m. intervals in all scatterplots. (**B**) Phosphoregulation of the eSAC phosphodomain and KNL1 analyzed by immunoblotting for the indicated proteins. (**C**) Left, Reversine treatment inactivates the SAC and significantly accelerates cell division (n=432 and 199 for untreated and Reversine-treated cells respectively, Mann-Whitney test). Right, Effect of Reversine on eSAC activity based on the partially Reversine-resistant Mps1S611R kinase domain (n=621 and 1193 for Rapamycin and Rapamycin + Reversine respectively; 2 trials). (**D**) Effect of the Aurora B inhibitor ZM447439 on the eSAC-induced mitotic arrest (n=931 and 205 for Rapamycin and Rapamycin + ZM447439 respectively; 2 trials). (**E**) Activity of the membrane-targeted eSAC (n=697 and 1056 for untreated and Rapamycin-treated cells respectively, 2 trials). Right, Confocal images display protein localizations as indicated. Scale bar ~ 5 μm.

To detect the involvement unknown cellular factors in the observed metaphase arrest, we tested whether the MELT motifs in the eSAC phosphodomain and the kinase activity of the eSAC kinase domain were necessary to induce the mitotic arrest. For this, we constructed two different cell lines: one expressed an analog-sensitive version of the eSAC kinase domain along with the eSAC phosphodomain; the other expressing wild-type eSAC kinase domain, but the non-phosphorylatable version of the eSAC phosphodomain (Sliedrecht et al., 2010; Vleugel et al., 2013). Inhibition of the analog-sensitive eSAC kinase domain or elimination the phosphorylation sites in the eSAC phosphodomain prevented the rapamycin-induced metaphase arrest (Figure S2A and S2B respectively). Thus, the catalytic activity of the eSAC kinase domain and the Mps1 phosphorylation sites within the eSAC phosphodomain are both required for the rapamycin-induced metaphase arrest. These data demonstrate that phosphorylation of the MELT motifs in KNL1 by Mps1 in the cytosol is sufficient to institute a metaphase arrest in human cells.

### eSAC activity is independent of kinetochores

We next sought to confirm that the observed eSAC activity is completely independent of the kinetochore-based SAC activation machinery. To assess this kinetochore-independence biochemically, we synchronized cells in G1/S, and then released them from the experimental block in media containing either Rapamycin to activate the eSAC or 100 nM nocodazole to depolymerize spindle microtubules and activate the SAC (Figure 2B). By probing cell lysates from these conditions with phospho-specific antibodies against the 13^th^ MELT motif in KNL1, we found that the two MELT motifs in the eSAC phosphodomain (containing a total of 6 MELT motifs) were phosphorylated only in the presence of rapamycin. In contrast, the eSAC MELT motifs were not phosphorylated if the SAC was activated by creating unattached kinetochores by treatment with nocodazole. Reciprocally, the MELT motifs in endogenous KNL1 were not appreciably phosphorylated in rapamycin-treated cells, but were strongly phosphorylated in nocodazole-treated cells (Figure 2A). Thus, there is negligible cross-talk in the phosphoregulation of the eSAC phosphodomain and endogenous KNL1.

Next, we sought to create an eSAC kinase domain that would remain active even when the endogenous Mps1k kinase was inhibited using Reversine to disable the kinetochore-based SAC activation pathway. To this end, we constructed an eSAC that utilizes a partially Reversine-resistant allele of the Mps1 kinase domain (Koch et al., 2015). When cells expressing this eSAC were treated with Reversine alone to inhibit endogenous Mps1 kinase activity, the duration of mitosis was significantly reduced in agreement with published data (scatterplot on the left in Figure 2C). However, when these cells were treated with Reversine and Rapamycin together, the duration of mitosis was comparable to the duration observed upon Rapamycin treatment alone (scatter plot on the right in Figure 2C). Reversine did partially reduce the mitotic delay caused by the eSAC as expected because the Mps1 allele used as the eSAC kinase domain is only partially Reversine resistant (IC_50_ ~ 130 nM versus ~ 30 nM for the wild-type Mps1 deduced from cell viability response to difference concentrations of the inhibitor; (Koch et al., 2015)).

We further tested the kinetochore-independence of the eSAC by assessing whether it requires Aurora B kinase activity. Aurora B is thought to contribute to SAC activity by promoting the recruitment of Mps1 and SAC proteins to kinetochores (Ditchfield et al., 2003; Santaguida et al., 2011; Saurin et al., 2011). If kinetochores contribute to the operation of the eSAC, then Aurora B kinase inhibition should negatively impact eSAC activity. However, we found that eSAC operation was marginally enhanced, when cells were treated with the Aurora B inhibitor ZM447439 (Figure 2C). This experiment further reveals that the kinetochore-based SAC activation pathway is not required for the eSAC-induced metaphase arrest.

As the final validation of kinetochore-independent operation of the eSAC, we targeted the eSAC phosphodomain to the plasma membrane by adding a palmitoylation sequence at its N-terminus (Figure 2D, top micrograph). The eSAC kinase domain remained cytosolic. When rapamycin was added to the growth media, the eSAC kinase domain re-localized to the membrane-tethered eSAC phosphodomain (Figure 2D, micrograph on the bottom). Importantly, this membrane-tethered eSAC activator induced a potent mitotic arrest that is similar in strength to the arrest induced by the cytosolic eSAC version (scatterplot in Figure 2D). Together, these experiments demonstrate that the eSAC operates independently of the kinetochore in instituting a mitotic arrest.

### eSAC delays anaphase onset by stimulating the SAC signaling cascade

We next sought to define the events that occur downstream from the dimerization of the eSAC phosphodomain and eSAC kinase domain. To this end, we immunoprecipitated the eSAC phosphodomain from cells arrested in mitosis, and performed mass spectrometry analysis to identify interacting proteins. When the eSAC phosphodomain was isolated from cells arrested in mitosis by treatments with the microtubule depolymerizing drug nocodazole, mass spectrometry analysis identified peptides from KNL1 and FKBP12, but not from Mps1 or any of the SAC proteins (Figure 3A). In contrast, when we affinity-purified the eSAC phosphodomain from cells treated with rapamycin, the same analysis identified the Mps1 kinase domain, Bub3, and Bub1. To identify dynamic interacting partners for the eSAC phosphodomain, we treated cells with the crosslinking agent formaldehyde prior to the affinity purification to trap weakly associated proteins. These purifications additionally isolated BubR1, a key component of the Mitotic Checkpoint Complex (MCC) (Sudakin et al., 2001). Thus, the eSAC phosphodomain recruits components of the SAC signaling cascade and the MCC only when it is phosphorylated by Mps1.

**Figure 3.**
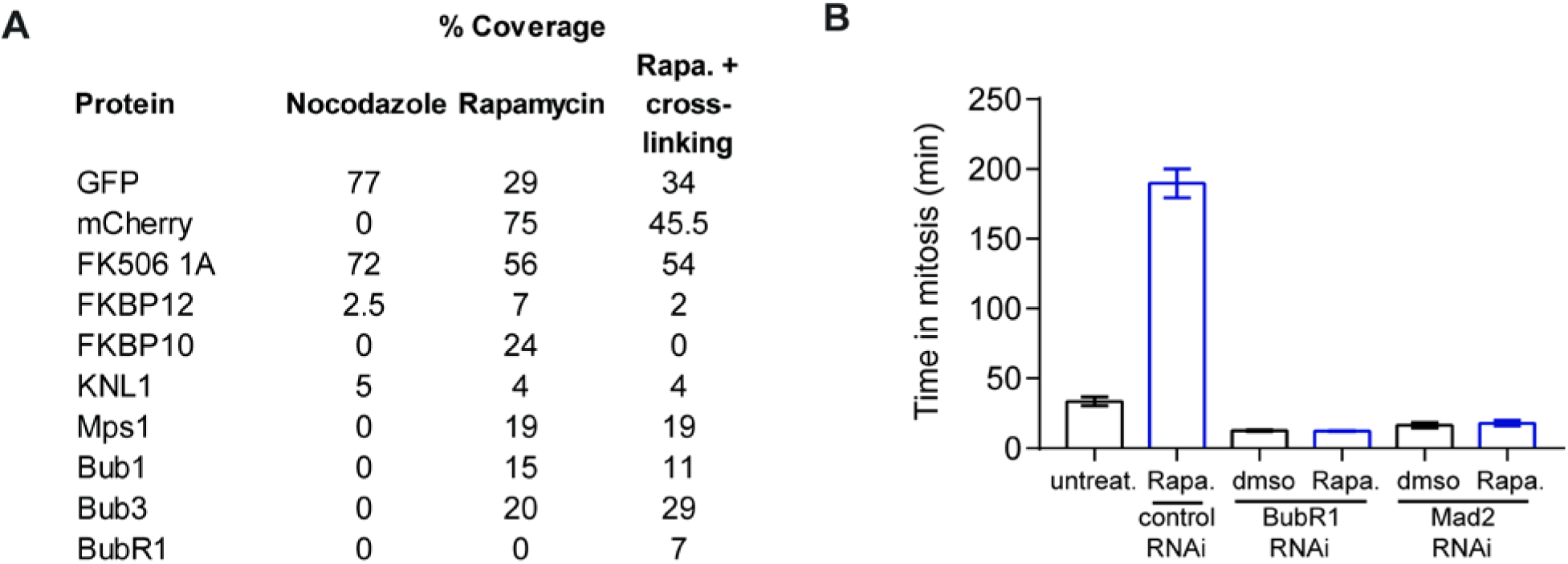
The eSAC phosphodomain interacts with SAC proteins and leads to the generation of the MCC. (**A**) Mass spectrometry analysis of immunoprecipitated eSAC phosphodomain under the indicated conditions. (**B**) Effect of RNAi-mediated depletion of either BubR1 or Mad2 on eSAC activity (n = 78, 390, 300,191, 72 and 140 respectively, 2 trials). Bar height = mean, error bars display s.e.m. Horizontal lines in scatter plots display mean +/- s.e.m.

Next, we tested whether the eSAC-induced metaphase arrest requires the formation of the MCC. We activated the eSAC in cells depleted for two of the essential MCC components: BubR1 or Mad2, using RNAi (Experimental Procedures). In both cases, rapamycin-treatment was unable to induce a mitotic arrest (Figure 3B). Instead, these cells underwent a significantly accelerated mitosis, consistent with the expected effect of depleting MCC components that are essential for arresting anaphase onset. These data indicate that the formation of the MCC is essential for the metaphase arrest induced by the eSAC.

Together, the results so far show that the eSAC generates phosphorylated MELT motifs in the cytosol, which then recruit SAC proteins and catalyze the formation of the MCC. The MCC then inhibits APC/C, and thus delays anaphase onset. Thus, the eSAC is a minimal, but potent system that institutes a controllable biochemical block to anaphase without interfering with the mechanics of cell division.

### Using the eSAC to determine the dose-response relationship for SAC signaling

The ability to hijack the SAC activation mechanism presented us with the unique opportunity of investigating the biochemical design and physiological operation of the SAC. Unlike the localized signaling activity of tightly clustered KNL1 molecules at unattached kinetochores, the eSAC activator complex is a diffusive, cytosolic complex of two proteins. Once formed, this complex is expected to be highly stable, because the binding affinity of the FKBP12 and FRB in the presence of rapamycin is high (*K_D_* ~ 10 nM, (Banaszynski et al., 2005)). The binding affinity of the eSAC kinase domain and the eSAC phosphodomain is likely to be much higher, because the eSAC uses two tandem repeats of the FKBP12 domain. The dosage of this activator complex is a key parameter. We expected that the concentration of the conditionally expressed eSAC kinase domain would be lower than that of the constitutively expressed eSAC phosphodomain, and hence it would be the factor that limits the formation of the eSAC activator complex. Using antibodies against the C-terminus of Mps1, we also confirmed that the abundance of the Mps1 kinase domain is similar to that of endogenous Mps1 kinase (Figure S3). These data indicate that eSAC activator complex concentration is comparable to the concentration of the endogenous, kinetochore-based KNL1-Mps1 activation machinery. Finally, the eSAC does not interfere with the mechanics of mitosis (Figure 2). Therefore, the eSAC should allow a quantitative determination of high-resolution ‘dose-response’ characteristics of the SAC signaling cascade: how the concentration of the stimulating signal controls the delay in anaphase onset.

We undertook a systematic determination of the dose-response characteristics using long-term imaging of cells expressing the two eSAC components. In this analysis, we used the 11^th^, 12^th^, 13^th^, and 14^th^ MELT motifs of KNL1 for which the kinetochore-based SAC signaling activity has been previously characterized (Vleugel et al., 2015; Vleugel et al., 2013). These MELT motifs show varying degrees of effectiveness in recruiting Bub3-Bub1 to unattached kinetochores (Figure 4A). It will become apparent in the following sections that this MELT motif property plays a key role is shaping the dose-response curves.

**Figure 4.**
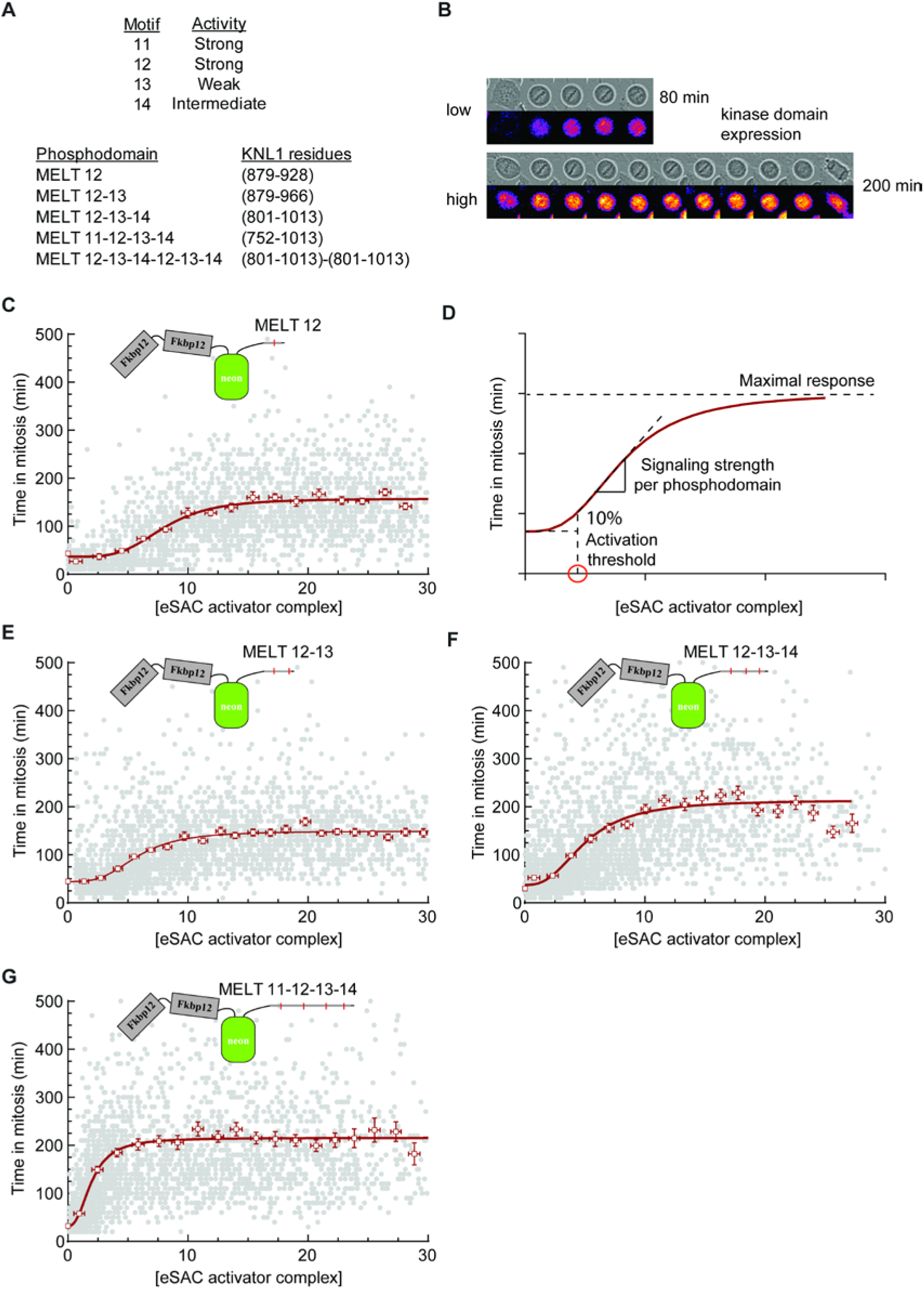
Dose-response curves for the SAC signaling cascade reveal properties of the KNL1 phosphodomain and characteristics of the signaling cascade. (**A**) Biochemical characteristic (top) and schematics (bottom) of the eSAC phosphodomains used in this study. (**B**) Montages of bright-field images and fluorescence heat-maps showing mitotic delay in cells expressing an eSAC phosphodomain with 1 MELT motif and different levels of the eSAC kinase domain (Δt = 20 minutes). (**C**) Dose (eSAC kinase domain fluorescence at the beginning of mitosis) vs. response (duration of mitosis) relationship for the eSAC phosphodomain with 1 MELT motif (n = 2572 from ≥ 2 trials). Each gray circle represents one cell. Open squares represent the means of binned data; error bars represent s.e.m. Red curve represent fit of the averages with a 4-parameter sigmoid. Duration of mitosis at 0 eSAC abundance was obtained from the mean duration of mitosis for the respective cell line in the absence of rapamycin. (**D**) Three characteristics of the eSAC phosphodomain and the SAC signaling cascade revealed by the dose-response curve (red curve – simulated dose-response curve). (**E-G**) Dose-response data for the indicated eSAC phosphodomains (n = 2969, 3043, and 2791 respectively from ≥ 2 trials). Red curves are 4-parameter sigmoid fits as before.

### Dose-response characteristics of the eSAC employing one MELT motif

We initiated our analysis of the SAC signaling cascade by using a version of the eSAC that employs just one MELT motif from KNL1 (Figure 4B). This eSAC activator complex represents the basic signaling unit in the SAC signaling cascade. In the experiments that follow, we exploited the induced expression of the eSAC kinase domain to obtain a cell population with highly heterogeneous levels of the eSAC kinase domain ranging from undetectable to relatively high levels (Supplemental Movies S4-S7). The abundance of the constitutively expressed eSAC phosphodomain displayed a much smaller cell-to-cell variance. Consequently, in the majority of cells, the abundance of the eSAC activator complex was limited by the abundance of the eSAC kinase domain, which was evidenced by the dependence of the observed mitotic delay on the cellular level of the eSAC kinase domain (Figure S4A-B). Therefore, to determine the dose-response characteristics, we used the average mCherry fluorescence in a given cell at the beginning of mitosis as the reporter for the eSAC activator complex. Using a custom automation script that uses the characteristic spherical shape of mitotic HeLa cells to detect and track them over time, we also determined the time that the cell spent in mitosis (see Experimental Procedures). This methodology enabled the analysis of thousands of cells and the extraction of dose-response characteristics for a given eSAC activator complex with high resolution.

We found that the cellular abundance of the eSAC activator complex affected the duration of mitosis systematically (Figure 4C). Strikingly, the dose-response curve displayed a sigmoidal trend (solid red line in Figure 4C). The sigmoidal dose-response curve reveals three physiologically significant features that quantitatively define the ability of the eSAC activator complex containing a single MELT motif to stimulate the SAC signaling cascade (Figure 4D). First, this analysis defines an activation threshold based on the abundance of the eSAC activator complex required to increase mitotic duration by 10% over the baseline duration (red circle in Fig. 4D). This threshold corresponds to the situation wherein one or a small number of unattached kinetochores activate the SAC. It relates to the minimum number of MELT motifs that will enable a single kinetochore to delay anaphase onset. Second, at concentrations beyond this activation threshold, the cellular response to increasing eSAC activator complex abundance was proportional. The steepness of the increasing response in this regime, obtained from the 4-parameter sigmoid fit, indicates the signaling strength per eSAC phosphodomain (dashed line in Figure 4D). This property, which is related to the activation threshold, characterizes the signaling strength of the KNL1 phosphodomain. Third, this analysis identifies the maximal delay that can be induced by the eSAC (Figure 4D). The maximal delay induced by high concentrations of eSAC activator complexes corresponds to the situation wherein multiple unattached kinetochores are engaged in checkpoint signaling. Thus, these three characteristics provide quantitative insights into the properties and features of the eSAC phosphodomain and the physiological operation of the SAC signaling cascade.

The high activation threshold and the relatively low maximal response induced by the eSAC phosphodomain containing a single MELT motif are both sub-optimal for SAC function. The high threshold implies that a large number of KNL1 molecules containing a single MELT motif would be required to generate a noticeable delay in cell cycle progression. The maximal delay (~ 157 minutes) is significantly lower than the average mitotic delay of ~ 500 minutes upon treatment of cells with taxol, or ~ 1500 minutes upon treatment of cells with nocodazole (Collin et al., 2013). Consequently, the single MELT-containing phosphodomain provides a very weak stimulation to the SAC signaling cascade. It should be noted that the observed asymptotic plateau suggests either that the abundance of the eSAC phosphodomain limits the formation of eSAC activator complexes, or that the abundance or activity of one or more SAC proteins prevents further increase in SAC signaling.

### Dose-response characteristics of eSAC phosphodomains employing up to 4 MELT motifs

To evaluate the effects of higher numbers of MELT motifs present in the KNL1 phosphodomain, we next investigated the dose-response relationship using eSAC phosphodomains with larger numbers of MELT motifs. We constructed three stable cell lines expressing eSAC phosphodomains containing two, three, and four MELT motifs respectively (Figure 4A), and determined their dose-response curves using the methodology described above (Figure 4E-G). The dose-response curve was sigmoidal in each case. Increasing numbers of MELT motifs per phosphodomain gradually reduced the activation threshold and increased the signaling strength of the phosphodomain (Figure 4E-G; see Figure 5D). The maximal, asymptotic delay in anaphase onset did not increase for the eSAC phosphodomain with 2 MELT motifs, but increased by ~ 40% for the eSAC phosphodomains containing 3 and 4 MELT motifs (Figure 4E-G; Figure 5E).

**Figure 5.**
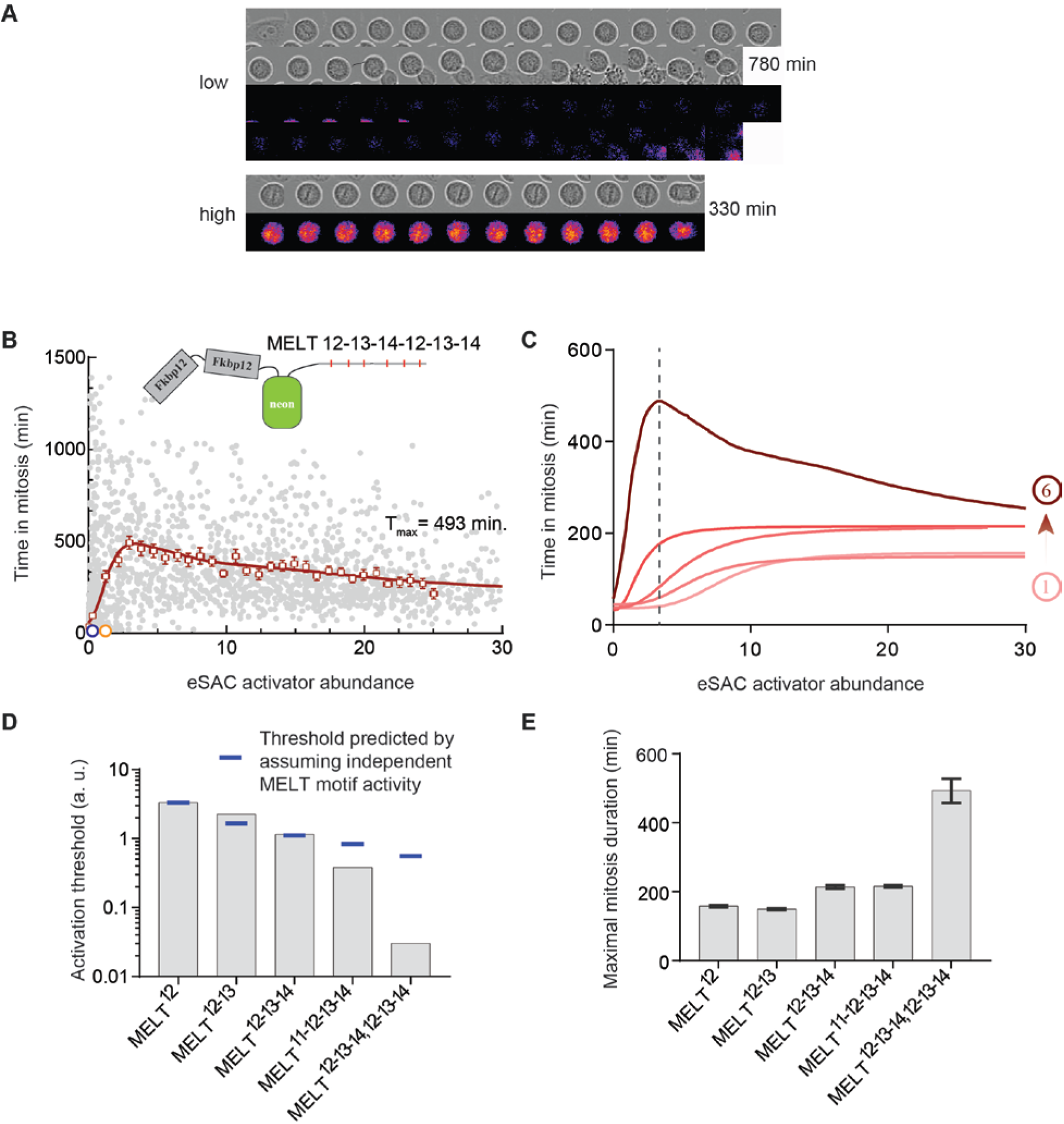
Emergence of synergistic signaling when multiple MELT motifs in the eSAC phosphodomain recruit SAC proteins. (**A**) Micrographs of two cells undergoing delayed mitosis due to the activity of an eSAC employing the phosphodomain harboring 6 MELT motifs (Δt = 30 minutes). (**B**) Dose-response curve for the eSAC phosphodomain with 6 MELT motifs. Each gray circle represents one cell. Open squares represent the means of binned data; error bars represent s.e.m. The red curve displays data smoothed with the Lowess filter (n = 2705 from ≥ 2 trials). (**C**) The average dose-response curves observed using the five eSAC phosphodomains. The dashed line separates the low and high dosage regimes for the eSAC employing 6 MELT motifs. (**D**) Activation thresholds and (**E**) maximal mitotic duration for the five eSAC phosphodomains. The values for each characteristic were either obtained from the sigmoidal fits or graphically by interpolation for the complex dose-response curve for the phosphodomain containing 6 MELT motifs.

These data show that the increased number of MELT motifs per phosphodomain proportionally reduce the activation threshold (Figure 5D). Thus, the additional MELT motifs expand the signaling capacity of the eSAC phosphodomain, and enable the eSAC to achieve strong stimulation of the SAC signaling cascade for the same molar concentration of the eSAC phosphodomain. Interestingly, the maximal delay in anaphase onset does not follow this pattern. It increases only for the eSAC phosphodomains with 3 and 4 MELT motifs (~ 40%, Figure 5E), and the observed increase is not proportional to the increase in the number of MELT motifs (~ 200%). As discussed above, the maximal delay could be either due to a mechanism that is intrinsic to the eSAC (saturating concentration of the eSAC activator complex) or extrinsic to it (limited abundance of SAC signaling proteins). If the formation of the eSAC activator complex is indeed the limiting factor, then the maximal delay in mitosis should increase in proportion with the number of MELT motifs per phosphodomain. This is because each eSAC phosphodomain provides a higher signaling capacity for the same molar concentration of the eSAC activator complex. The data clearly show that this is not the case. This observation strongly suggests that the limited abundance of one or more SAC proteins restricts eSAC activity at high concentrations of the eSAC activator complex (Aravamudhan et al., 2016; Heinrich et al., 2013).

### Dose-response characteristics of the eSAC phosphodomain employing 6 MELT motifs

The data above demonstrate that higher numbers of MELT motifs in the eSAC phosphodomains generate a nearly proportional increase in its signaling strength (Figure 5D). This trend predicts that eSAC phosphodomains with even larger numbers of MELT motifs will have a correspondingly smaller activation threshold. However, they likely will not achieve longer, maximal mitotic delay, because of the limited availability of SAC proteins. To test these predictions, we determined the dose-response relationship for an eSAC phosphodomain containing 6 MELT motifs. Importantly, this phosphodomain was created by duplicating the eSAC phosphodomain with 3 MELT motifs so that the biochemical characteristics of the MELT motifs used did not change.

The dose-response relationship for the eSAC phosphodomain with 6 MELT motifs was surprisingly complex, and it revealed a novel aspect of the biochemical design of the SAC (Figure 5A-B). This eSAC phosphodomain displayed a striking low activation threshold and high signaling strength, and it also achieved a disproportionately large increase in the maximal duration of mitosis (Figure 5B-E). Interestingly, the response gradually decreased as the dosage of the eSAC activator complex increased (dashed line in Figure 5B). Effectively, the eSAC activator complex containing the 6 MELT motif phosphodomain stimulated the SAC signaling cascade in a switch-like manner (Figure 5B). Importantly, the complex dose-response characteristics of this eSAC activator complex did not change even when it was tethered to the membrane or when the endogenous Mps1 kinase activity was inhibited (Figure S4C and D respectively). These data indicate that the observed response is an intrinsic property of the eSAC phosphodomain.

The disproportionately large increase in signaling strength generated by the 6 MELT eSAC phosphodomain is indicative of synergistic activity. We propose that the synergism emerges from the simultaneous recruitment of SAC proteins by more than one MELT motif in the same phosphodomain. This allows the eSAC phosphodomain to generate a signal that is significantly larger than the additive signaling activity supported by the individual MELT motifs present in the phosphodomain. This conclusion is further bolstered by the gradual decline in the response observed with increasing concentrations of the eSAC activator complex (Figure 5B-C). As the eSAC activator complex concentration increases, these complexes compete with one-another to recruit SAC proteins from the limited pool available in the cell. Consequently, individual eSAC activator complexes can no longer recruit multiple SAC proteins, and hence their synergistic activity weakens. Therefore, at high eSAC concentration the dose-response curve for the phosphodomain with 6 MELT motifs approaches the asymptotic plateau for eSAC phosphodomains containing 3 or 4 motifs (Figure 5C).

### A mechanistic model of the stimulation of the SAC signaling cascade by the eSAC explains the observed dose-response characteristics

To understand the mechanistic basis of the dose-response curves, we next devised a two-component mathematical model. This model simulates the stimulation of the SAC signaling cascade by the eSAC, and then calculates the effect of this stimulation on the timing of anaphase onset (Figure 6; Supplemental Information). Our model is based on the following biological data. Each MELT motif in KNL1 displays an apparent binding affinity for Bub3-Bub1 as revealed by their effectiveness in recruiting Bub3-Bub1 to unattached kinetochores (Vleugel et al., 2015). Furthermore, the Bub3-Bub1 recruitment activity of individual MELT motifs appears to be additive, which suggests that they act independently (Vleugel et al., 2015). The duration of mitotic arrest correlates with the number of SAC proteins recruited by KNL1 at kinetochores, and with the steady-state MCC concentration in the cell (Collin et al., 2013; Dick and Gerlich, 2013; Heinrich et al., 2013). Finally, we have shown that the eSAC activator complex recruits SAC proteins including Bub3, Bub1, and BubR1 to generate the MCC (Figure 3), and that the concentration of one or more SAC proteins is lower than the maximal dosage of the eSAC activator complex (Figures 4-5).

**Figure 6.**
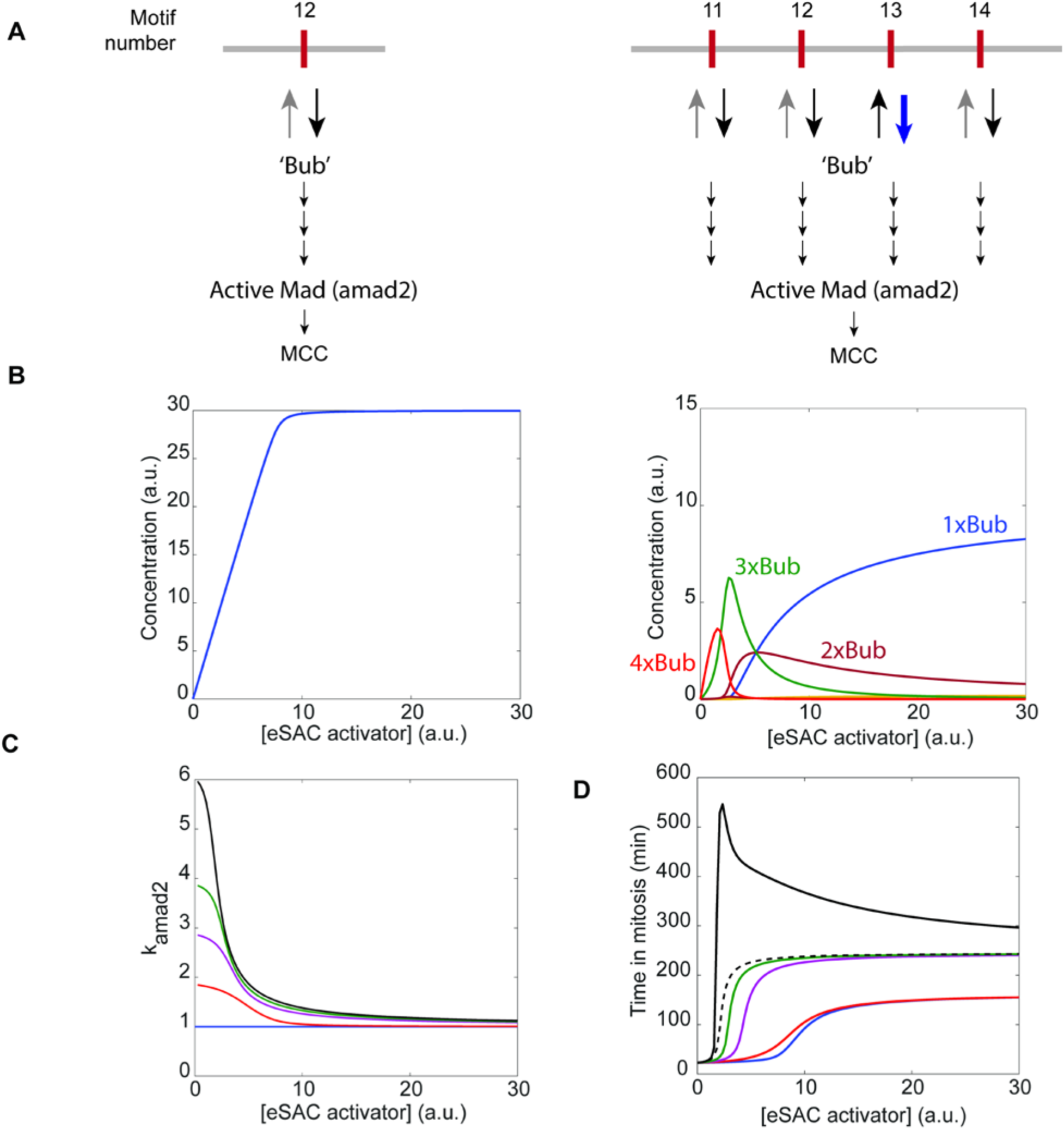
Mathematical model of the eSAC and simulation of the dose-response curves. (**A**) Model of eSAC phosphodomains containing 1 and 4 MELT motifs respectively. It represents the binding of all SAC proteins by the single “Bub” unit. The amino acid sequence of each MELT motif (labeled by its motif number in KNL1) determines its binding affinity for Bub. (**B**) Graphs display the steady-state abundance of different Bub-bound phosphodomain species for eSAC phosphodomains containing one (left) and four (right) MELT motifs. Each species is designated by the number of Bub proteins recruited by the respective phosphodomain. (**C**) The rates of the active or closed-form Mad2 generation (k_amad2_) supported by the five eSAC versions. To simulate the metaphase to anaphase transition, the second part of the model assumes that the rate of MCC generation is proportional to k_amad2_. (**D**) Simulated dose-response curves obtained by relaying the signaling activity of the eSAC model to a previously described model for a bistable switch that governs mitotic exit (He et al., 2011). The dashed and solid black curves display the dose-response characteristics for the phosphodomain with 6 MELT motifs without and with synergistic activity respectively.

Our model assumes that each MELT motif in the eSAC activator complex recruits SAC proteins with a characteristic affinity (Figure 6A, Table S1). We represent all the different SAC proteins by a single protein species named ‘Bub’ (Figure 6A). This simplification of the SAC signaling cascade is necessary, because quantitative measurements for the recruitment reactions are unavailable. Moreover, the biochemical interactions that recruit SAC proteins to the eSAC activator complexes are shared by all five versions of the eSAC phosphodomain. Therefore, this simplification does not detract from the main goal of this simulation, which is to understand the mechanisms that give rise to the observed dose-response curves. Under this simplified scheme, we used the apparent binding affinity of each MELT motif and Bub3-Bub1 to simulate the interaction between the MELT motifs in the eSAC activator complex and the Bub protein. Therefore, the model calculates the steady state concentration of all possible species of the eSAC phosphodomain that are characterized by the MELT motifs recruiting Bub according to the mass action law (concentrations for eSAC phosphodomains with 1 and 4 MELT motifs displayed in Figure 6B). The model also assumes that the cellular abundance of the ‘Bub’ protein is lower than the maximum dosage of the eSAC activator complex. Due to this limitation on available Bub, the abundance of eSAC phosphodomains bound with different numbers of Bub strongly depends on the eSAC activator abundance (Figure 6B, Figure S5). Finally, the rate of generation of the active form of Mad2, and hence that of MCC, depends on the number of Bubs recruited by the eSAC phosphodomains (Figure 6C).

To simulate the response of the cell to the MCC generated by the eSAC, it was necessary to model anaphase onset. For this, we used our published model that employs a mathematical representation of a bi-stable switch to describe the irreversible metaphase to anaphase transition (Figure S6A) (He et al., 2011). The state of this switch is determined by the balance of the kinase activity of Cyclin B/CDK1 and an antagonizing phosphatase, and degradative activity of the APC/C and the antagonizing MCC (Figure S6A, also see Supplemental Information for details). We relayed the cumulative MCC generated by the eSAC to this model to simulate the timing of anaphase onset, and thus obtained the ‘time in mitosis’ as a function of the eSAC activator complex (Figure S6B-D).

This model captured the average dose-response characteristics of eSAC phosphodomains containing up to 4 MELT motifs (compare Figures 5C and 6D). However, this simple scheme did not reproduce the complex dose-response relationship for the phosphodomain with 6 MELT motifs (Figure 6D, dashed black curve). In this case, it was necessary to assume that eSAC phosphodomains that had more than one MELT motif bound by Bub produce MCC at a modestly higher rate (≤ 20% increase due to synergistic output, see Table S2). With this modification, the model accurately captured the dose-response characteristics for the eSAC phosphodomain containing 6 MELT motifs (Figure 6D). Our analysis demonstrates that the recruitment of SAC proteins to the phosphorylated MELT motifs in the eSAC activator complex governed by the mass action law, and followed by the formation of MCC explains can explain the observed dose-response data for the eSAC phosphodomains. It also shows that synergistic signaling activity of eSAC phosphodomains that engage multiple MELT motifs in SAC signaling explains the complex dose-response data.

## Discussion

### The SAC: a biochemical approximation of the toggle-switch

The complexity of the SAC signaling cascade and its localized activity to discrete, nanoscopic sites in the cell make its physiological operation inaccessible to systematic and quantitative biochemical analyses. Consequently, fundamental questions regarding the biochemical design and physiological operation of the SAC remained unanswered. Our analysis of the eSAC as a soluble, kinetochore-independent SAC activator, provides an unprecedented view of the physiological operation of the SAC signaling cascade, and reveals novel features of its biochemical design. We find that the biochemical signaling cascade of the SAC operates as a range-limited rheostat. However, the KNL1 phosphodomain maximally stimulates this cascade using a synergistic signaling activity that involves multiple proximal MELT motifs, and elicits a switch-like response.

### Limited abundance of SAC proteins restricts SAC signaling activity

We used the eSAC to incrementally stimulate the SAC signaling cascade, and deduce its dose-response characteristics with high resolution. The comparative analyses of eSAC phosphodomains harboring increasing numbers of MELT motifs quantitatively reveal how larger numbers of MELT motifs strengthen the signaling output of KNL1. Effects of the differential Bub3-Bub1 recruitment activity of individual MELT motifs (listed in Figure 4A) are also apparent in these data (Vleugel et al., 2015). The maximal mitotic delay generated is the same for eSAC phosphodomains harboring the 11^th^ (strong), and the 11^th^ and 12^th^ (weak) MELT motifs, but it increases when additional stronger MELT motifs are added to the eSAC phosphodomain (Figures 5E). A mechanism involving ‘Bub’ interaction with the MELT motifs governed by mass action in our simplified mathematical model of the SAC explains how stronger MELT motifs can maintain higher steady-state levels of MCC (Figure S5, Figure 6D).

Importantly, our quantitative analyses provide several key insights into the physiological operation of the SAC. Under conditions of low to moderate stimulation, the SAC signaling cascade produces a proportional response in the form of delayed anaphase onset. Effectively, this behavior of the SAC resembles that of a rheostat resisting anaphase onset (Collin et al., 2013). This behavior also provides clear evidence that the anaphase-inhibitory signal produced by unattached kinetochores is not amplified in the cytoplasm (Mariani et al., 2012; Sear and Howard, 2006; Simonetta et al., 2009). The saturation of the maximal mitotic delay with increasing concentrations of the eSAC activator complex is also a highly significant aspect of the operation of the SAC signaling cascade. The inability of the eSAC to further stimulate the SAC signaling cascade reveals that the limited abundance of SAC proteins restrains SAC signal generation. This mechanism can be expected to similarly limit SAC signal generation when large numbers of unattached kinetochores participate in SAC signaling.

### Synergistic signaling by the KNL1 phosphodomain produces switch-like SAC activation

The proportional response of the SAC to increasing concentrations of eSAC activator complexes containing up to 4 MELT motifs implies that the kinetochore should contain a very large number of MELT motifs to ensure that a single kinetochore can generate a robust quantity of MCC. Indeed, human KNL1 contains 19 MELT motifs (Vleugel et al., 2013). However, ~ 5 of these MELT motifs possess degenerate amino acid sequences, and only 6 out of the remaining motifs participate in Bub3-Bub1 recruitment in nocodazole-treated cells (Tromer et al., 2015; Vleugel et al., 2015). These puzzling observations regarding KNL1 activity can be explained by the emergence of synergistic activity when multiple MELT repeats in a KNL1 phosphodomain engage in SAC signaling.

Very low dosage of the eSAC activator complex containing 6 MELT motifs generate a strong, increasing response from the SAC signaling cascade, but this response decreases with further increase in the dosage of the eSAC activator complex (on either side of the dashed line in Figure 5C). It is important to note that the independent Bub3-Bub1 recruitment activity of MELT motifs in the eSAC activator complexes implies that the total amount of Bub3-Bub1 recruited does not decrease with increasing dosage of any of the eSAC activator complexes. Therefore, the disproportionate increase and then gradual decay in the response of the SAC signaling cascade is best explained by the emergence of synergistic activity when multiple MELT motifs in one eSAC phosphodomain recruit SAC proteins simultaneously. The synergistic activity will enhance the rate of MCC generation, and hence produce a longer than expected delay in anaphase onset. As the concentration of the eSAC activator complex increases, the fraction of phosphodomains that can bind multiple SAC proteins drops, the synergistic signaling activity is reduced, and hence the response of the cell asymptotically decays to the level achieved by eSAC phosphodomains harboring lower numbers of MELT motifs (Figure 5C, Figure 6D).

### Automatic gain modulation of the signaling from individual kinetochores enables switch-like response of the SAC

The characteristics of the KNL1 phosphodomain and the SAC signaling cascade discussed above suggest an elegant model for how the SAC realizes a switch-like response to the presence of unattached kinetochores (Figure 7). We propose that the synergistic activity of KNL1 combined with the limited abundance of one or more SAC proteins in the cell together institute automatic gain control on the signaling strength of individual kinetochores. When multiple unattached kinetochores participate in SAC signaling, they compete with one another in recruiting from the limited pool of SAC proteins (Figure 7B). This competition limits the signal that can be generated by each kinetochore by frustrating synergistic activity of the KNL1 phosphodomain. However, because the number of signaling kinetochores in the cell is high, the cumulative MCC level remains high, and anaphase is significantly delayed (Figure 7C, red curve). As the number of signaling kinetochores in the cell drops, the abundance of SAC proteins relative to the signaling kinetochore number increases (Figure 7B, blue curve). As a result, KNL1 molecules in these kinetochores can now recruit multiple SAC proteins and realize synergistic signaling (depicted as a threshold event by the dashed line in Figure 7B and C). The synergistic signaling by KNL1 molecules increases their signaling output non-linearly. Consequently, the MCC levels in the cell do not drop in proportion to the number of unattached kinetochores. Such automatic gain control on the signaling strength of unattached kinetochores will allow the SAC to maximize chromosome segregation accuracy and minimize the duration of mitosis.

**Figure 7.**
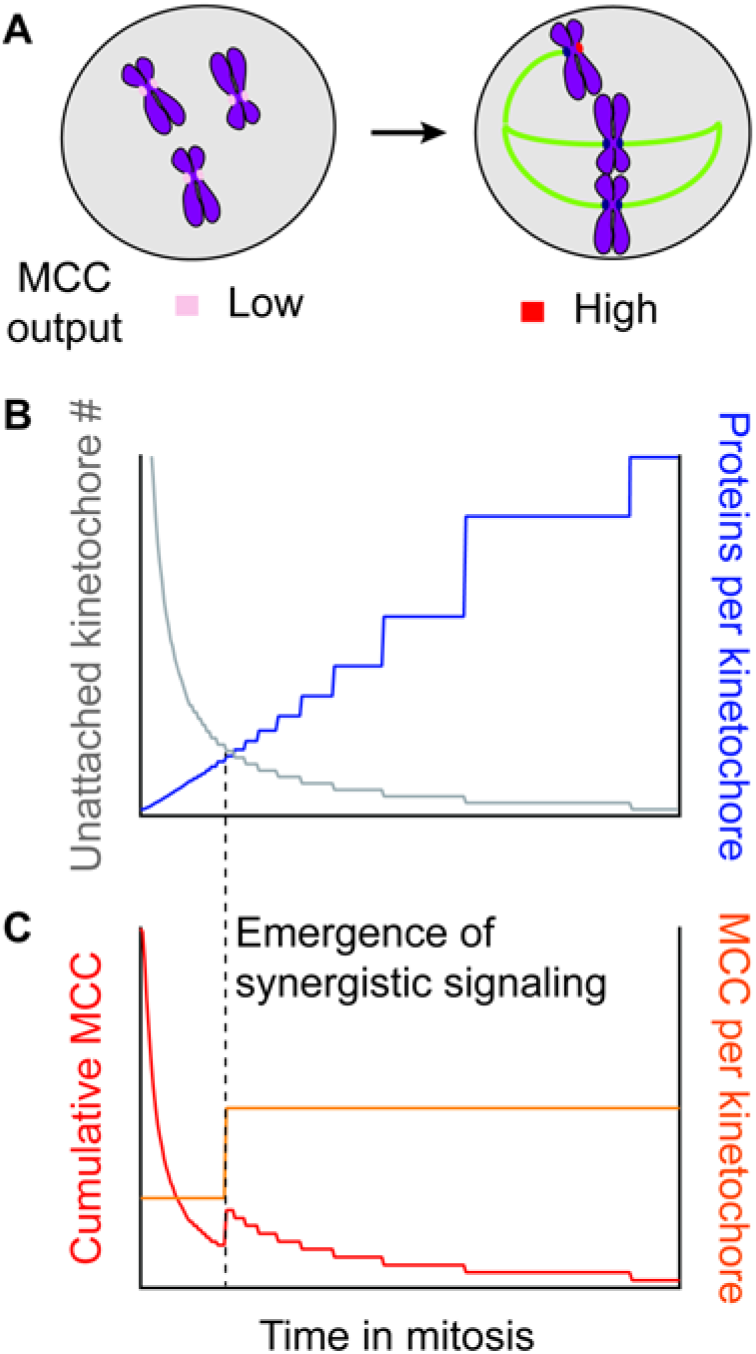
Hypothesis to explain the mechanisms that enact a switch-like SAC. **(A)** The SAC signal output (MCC generation) per kinetochore increases with the falling number of unattached kinetochores in the dividing cell. (B) When the number of unattached kinetochores is high, the number of available SAC proteins per kinetochore is low (blue curve). This restricts the MCC output from individual kinetochores, but the cumulative MCC generation is high because the number of signaling kinetochores is high. The availability of SAC proteins increases as the number of unattached kinetochores decreases (gray curve). (C) Increased availability of SAC proteins facilitates synergistic signaling (depicted in this cartoon as a threshold event by the dashed line) from the small number of unattached kinetochores, which increases the signal output per kinetochore (orange curve). Consequently, the cumulative MCC generation (red curve) does not track linearly with the number of unattached kinetochores.

### A versatile, genetically encoded tool for controlling the duration of mitosis

In addition to revealing the biochemical design and physiological operation of the SAC, the eSAC will find use in other important investigations of the role of the SAC in cell and developmental biology. It provides an excellent tool as a potentiometer for measuring the inherent signaling potential of the SAC signaling cascade, and detecting subtle defects in its potency. This application is especially relevant for studying the SAC in cancer cell biology. Cancer cells are thought to have a defective SAC, because they express many SAC proteins aberrantly (Holland and Cleveland, 2009). However, the significance of the aberrant protein expression to tumorigenesis or tumor cell proliferation is unclear, because the SAC itself is operational as assayed by challenging cancer cells with nocodazole (Tighe et al., 2001). However, it is important to note that nocodazole-mediated activation of the SAC reveals only the maximum signaling strength of the SAC, because it creates many unattached kinetochores. It cannot detect small changes in its potency that will be critical for a small number of unattached kinetochores to be able to activate the SAC. The eSAC will be ideal for detecting such subtle changes. The eSAC also provides a genetically encoded tool for controlling the duration of mitosis in cell and developmental biology. It is based on the highly conserved domains of two mitotic proteins that are unlikely to have functions outside of mitosis. Therefore, the eSAC can be used establish a genetically-encoded, generalizable, and tunable control over the onset of anaphase, and hence the length of mitosis, in cell and developmental biological studies.

## Acknowledgments

The authors would like to thank Pavithra Aravamudhan, Mara Duncan, Michael Lampson, and Yukiko Yamashita for a critical reading of the manuscript. This work was supported by grants from the NIH/National Institute of General Medical Sciences: GM088313 to IMC, GM078989 to JJT (subcontracted from Colorado State University) and GM112992 to APJ, and by a Scholar award to IMC from the Leukemia & Lymphoma Society. Use of the Incucyte system was made possible by a generous gift from the Richard Tam Foundation to Prof. Sue O’Shea (Cell & Developmental Biology, University of Michigan Medical School). The authors also thank Dr. Eugene Makeyev for providing HeLa acceptor cell line, Dr. Michael Lampson for the generous gift of Mps1 plasmids, and Dr. Geert Kops for the generous gift of plasmids and anti-phospho-MELT antibody.

## Author contributions

APJ and IMC designed the experiments. APJ and CC generated cell lines and conducted Incucyte experiments. CC wrote the program to automate image analysis. APJ and AF analyzed the data. AF conducted Western blot analysis. IPW and DMK performed live cell and immunofluorescence imaging experiments. IW performed mass spectrometry analysis. AB and JJT constructed the mathematical model and performed simulations. APJ, IMC, and JJT wrote the manuscript.

## Method Details

### Generation of stable HeLa cell lines

Each cell line used in the study was generated by integrating a bi-cistronic cassette at an engineered *Loxp* site in the HeLa genome via Cre-mediated recombination using plasmids and protocols developed and generously provided by the Lampson lab (Ballister et al., 2014; Khandelia et al., 2011). Several colonies of transformed cells obtained from each transfection were pooled together, screened for the expression of integrated genes, and then cultured as necessary for experimentation.

### Cell culture and synchronization

Cells were maintained in DMEM with 10% FBS, 1% Pen/Strep at 37°C with 5% CO2, with 1 μg/ml Puromycin added to select for transformed cells. Expression of the Mps1 kinase domain was induced ~ 48 hours prior to the start of each experiment by adding Doxycycline to the culture medium (2 μg/ml final concentration achieved from a 1 μg/ml stock in DMSO). To synchronize at the beginning of S phase, asynchronous populations were treated with 2.5 mM Thymidine for 16 hours, released from the thymidine block for 9 hours, treated again with 2.5 mM Thymidine for 16 hours, and finally released from the second thymidine block.

### Chemicals

To induce the dimerization of Fkbp12 and Frb-labelled proteins, Rapamycin was added to the media ~ 1 hour prior to the start of each experiment (final concentration of 500 nM from a 500 μM stock in DMSO). To arrest cells in mitosis, Nocodazole was added to the media to a 100 nM final concentration from a 100 μM stock in DMSO. To inhibit Mps1 kinase activity, Reversine was added to the media to a final concentration of 500 nM from a 500 μM stock in DMSO. To inhibit Aurora B kinase activity, ZM-447439 was added to a final concentration of 10 μM from a 10 mM stock in DMSO.

### Transfection with siRNA

The following oligonucleotides were transfected by treatment with Lipofectamine3000 (Invitrogen) as per the manufacturer’s protocol: MAD2 10 nM (GAGUUCUUCUCAUUCGGCAUCAACA); BUBR1 10 nM (GAUGGUGAAUUGUGGAAUA[dT][dT]; (Bolanos-Garcia et al., 2011)), Negative control: ALLSTAR 20 nM.

### Live cell imaging

Before imaging, cells were shifted to Fluorobrite media supplemented with 10% FBS and 1% P/S). High resolution time-lapse imaging was conducted on a Nikon Ti-U inverted microscope equipped with a 100x, 1.4 NA oil immersion objective, Lumencor light engine for fluorescence excitation, and an Andor iXon DV897 EM-CCD camera. An environmental chamber (Chamlide TC, Quorum Inc.) was used to maintain optimal conditions for cell growth. MetaMorph 7.6 was used to drive the microscope and to run multi-position time-lapse experiments. In these experiments, cells grown on glass-bottom dishes were treated 500 nM rapamycin for 1 hour prior to imaging. Cells with a visible mCherry signal were selected for time-lapse imaging. During the imaging session, a bright-field, neonGreen (GFP), and mCherry image was acquired every 2 minutes for a total duration of ~ 2 hours.

Long-term imaging of the eSAC cell lines was conducted using the Incucyte Zoom Live Cell Imaging system (Essen Bioscience Inc.) equipped with a 20x phase objective. To conduct several experiments in parallel, cells were seeded in 12-well tissue culture plates (Corning) ~ 48 hours prior in media containing 2 μg/ml Doxycycline. One well in each experiment was left empty, and used to record background fluorescence at each time point during the experiment. Approximately 30-60 minutes prior to imaging, the growth media were exchanged with Fluorobrite media alone or media containing Rapamycin and small molecule inhibitors. In each well, we acquired a phase and fluorescence image every 10 minutes at four pre-selected positions. mCherry fluorescence was acquired using 900 ms exposure, while neonGreen fluorescence was recorded using a 300 ms exposure. The duration of each experiment was limited to ~ 24 hours in order to minimize the inclusion of cells that enter second mitosis during the course of the experiment.

### Image analysis

In imaging experiments conducted on the high-resolution microscope, we manually scored the duration of mitosis. In long-term imaging experiments, we partially automated this scoring using a custom script implemented in MatLab. Each experiment generated either two (phase and mCherry) or three (phase, GFP, and mCherry) image stacks. These image stacks were first pre-processed to remove stage jitter and background fluorescence. To remove stage jitter, phase images were registered as follows. First, we applied a simple threshold to the phase image stack to remove background, and then used median filter to minimize noise. Each phase image (with the exception of the very first image) was then registered with the preceding image in the stack using the MatLab function ‘imregtform’ using only translation to achieve registration. The transform function computed for this registration was applied to the corresponding image in the GFP and mCherry stack. Background fluorescence was removed from the GFP and mCherry image stacks, prior to registration, by subtracting from these stacks the average background fluorescence image in the respective channel obtained from the unseeded well in each experiment. The processed image stacks were then combined to create a composite image stack, which was used for tracking mitotic cells. Mitotic HeLa cells assume a characteristic spherical shape (average diameter ~ 25 μm). In phase images, these cells appear as circles defined by a high-contrast edge. To automatically detect these cells, we convolved each phase image with the cropped image kernel of a prototypical mitotic HeLa cell (constructed by 2-D averaging of phase contrast images of 19 manually selected mitotic HeLa cells). To remove non-mitotic features from the convolution, we applied a threshold to the convolution image, and then recorded the peaks in convolution as the centroids of circular features i.e. mitotic cells. The centroids appearing in successive frames were linked as belonging to the same cell, if they were no farther than 0.4 x cell diameter from one another. Gaps in linking were filled in, if a centroid persisted within the defined area over three successive time points. Cells that did not enter anaphase at the end of the imaging session were not considered for further analysis. Finally, annotated crops of the linked cell images were presented to the user as a montage in a graphical user interface (GUI). This GUI suppressed cells that divided in 10 minutes (one frame), except for experiments (Mad2 and BubR1 RNAi and Reversine treatment) wherein impaired SAC function significantly reduced the duration of mitosis. With this GUI, the user can either accept or reject the displayed cell montage, and if necessary, adjust the time frame of mitotic entry and exit based on the morphology of the cell as seen in the phase contrast image. A small fraction of the cells that experienced a long mitotic arrest and underwent cell death were also scored. Taxol treated cells did not undergo cytokinesis. They either underwent cell death or re-spread on the chamber surface. For these cells the duration of mitosis was scored as the time for which the cell maintained circular, mitotic appearance. After the final user input, the GUI calculated the duration of mitosis as the duration for which the circular morphology of the cell persists. It also recorded the mCherry and GFP fluorescence from the respective images by averaging the fluorescence intensity within a ~ 15 μm circle with its center on the centroid in the first image of the mitotic cell.

### Immunofluorescence

Immunofluorescence was performed as described previously (Kline et al., 2006). DNA was visualized using 10 μg/ml Hoechst. Immunofluorescence and live cell images were acquired on a DeltaVision Core deconvolution microscope (Applied Precision) equipped with a CoolSnap HQ2 CCD camera and deconvolved where appropriate. For immunofluorescence, approximately 10–20 Z-sections were acquired at 0.2 μm steps using a 100×, 1.4 Numerical Aperture (NA) Olympus U-PlanApo objective. Live cell imaging was performed using a 60×/1.42 NA Olympus U-PlanApo objective.

### Immunoprecipitations and Mass Spectrometry

The GFP tagged KNL1 phosphodomain was isolated from HeLa cells as described previously (Cheeseman and Desai, 2005). Co-purifying proteins were identified using a LTQ XL Ion trap mass spectrometer (Thermo Fisher Scientific) using MudPIT and SEQUEST software as described previously (Washburn et al., 2001). For the crosslinking IP experiments, cell pellets were resuspended with 1.2% formaldehyde in PBS and rocked gently for 10 min. After pelleting at 1,000 g, the cell pellet was quenched with 0.125 M glycine in PBS for 10 min. The cells were processed by sonication and treatment with detergent to solubilize cross-linked cell material, and then purified as for the noncrosslinked sample. The reversal of the crosslinks reversal was performed at 95°C for 5 min before tryptic digestion.

### Mathematical model of the eSAC and simulation of mitotic exit

The goal of this model was to elucidate the mechanisms that shape the average dose-response curves determined *in vivo*. The model consists of two components.

1. First, we calculate the steady-state concentrations of different species of a give eSAC phosphodomain defined by the interaction of SAC proteins with the MELT motifs in the phosphodomain. These interactions are governed by mass action, the concentration of SAC proteins, and the affinity of each MELT motif to SAC proteins. The SAC signaling cascade is highly complex, based on the recruitment of a large number of SAC proteins to the KNL1 phosphodomain (Figure 1B). Furthermore, quantitative measurements for this recruitment are not available. Therefore, significant simplification of this cascade was necessary in our simulation. To this end, we represented all of the SAC proteins by a single unit called Bub (Figure 6A). This simplification does not affect our main goal, because the SAC protein recruitment mechanisms are the same for all five eSAC phosphodomains. Next, the model uses the steady-state concentrations of eSAC phosphodomain species to calculate the rate of closed/active Mad2 generation. Following *in vivo* data, the model assumes that the rate of closed/active Mad2 generation is proportional to the number of SAC proteins interacting with the eSAC phosphodomain (Collin et al., 2013).
2. The second component of our model is designed to simulate the observed response of the eSAC cells, which is the duration of mitosis (Figure 4 and 5). The eSAC activity affects the duration of mitosis in the following manner. The rate of closed/active Mad2 generation calculated in the first component determines the steady-state concentration of MCC. The activity of the Anaphase Promoting Complex/Cyclosome is inversely proportional to the steady-state MCC concentration (Collin et al., 2013; Heinrich et al., 2013). This activity decides the rate of Cyclin B degradation. To simulate mitotic exit, these calculations were relayed to a mathematical representation of a bistable switch (He et al., 2011). In this model, once Cyclin B level in the cell falls below a threshold value, two positive feedback loops commit the cell to mitotic exit.

The individual steps of our two-component model are discussed in detail below.

#### Simulation of SAC protein recruitment by the eSAC phosphodomain

The eSAC is activated when the kinase domain of Mps1, which is fused with FRB is dimerized with the eSAC phosphodomain containing the specified number of MELT motifs, which is fused to 2xFKBP12. Binding of FKBP12-FRB induced by rapamycin is a high affinity reaction (*K_D_* ~ 10 nM) (Banaszynski et al., 2005). The use of two FKBP12 motifs will further increase this affinity significantly. Therefore, the Mps1-eSAC phosphodomain complex, i.e. the eSAC activator, is likely to be stable over the course of the experiment. We assume that Mps1 can phosphorylate each MELT motif in the unstructured phosphodomain with equal effectiveness. This assumption, together with the stability of the Mps1-eSAC phosphodomain complex, implies that all the MELT motifs will be phosphorylated in the presence of rapamycin.

A previous study found that the biochemical activity of each MELT motif depends on its amino acid sequence (Vleugel et al., 2015). Although this study did not measure the dissociation constants for each MELT motif, it quantified the effectiveness with which each MELT motif recruits Bub3-Bub1 to unattached kinetochores (Figure 4A). To reflect this in the model, we assigned the same rate of binding (*k_f_*, Table S1) of the ‘Bub’ protein for all MELT motifs, but assigned a much larger unbinding rate (*k_r_*, Table S1) for the weakest MELT motif (MELT 13). Furthermore, the model assumes that the Bub protein interacts with each MELT motif in the eSAC phosphodomain independently of the other MELT motifs, if they are present.

At each eSAC kinase domain concentration value (symbolized as [Mps1]), the eSAC phosphodomains in a cell can assume different Bub binding states. For example, for phosphodomains with two MELT motifs the possible states are M_12,13_, M_12B,13_, M_12,13B_, and M_12B,13B_, where the number in the subscript followed by the letter ‘B’ means that the particular MELT motif under consideration is bound to Bub. The time evolution of concentration of different Bub binding states is given by:
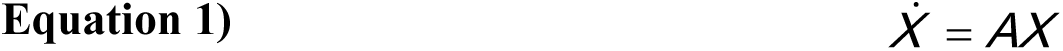

In this equation, *X* is a column vector of the concentrations {*x*_1_, *x*_2_, …, *x_N_*} of the *N* different Bub-binding states of the phosphodomain, and *A* is a matrix of rate constants discussed above. The concentrations satisfy the constraint:
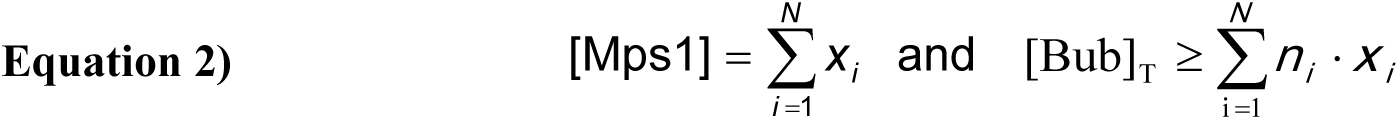

Here, *n_i_* = number of Bub units bound to species *x_i_*. For example, for a phosphodomain with two MELT motifs, *X* and *A* are:
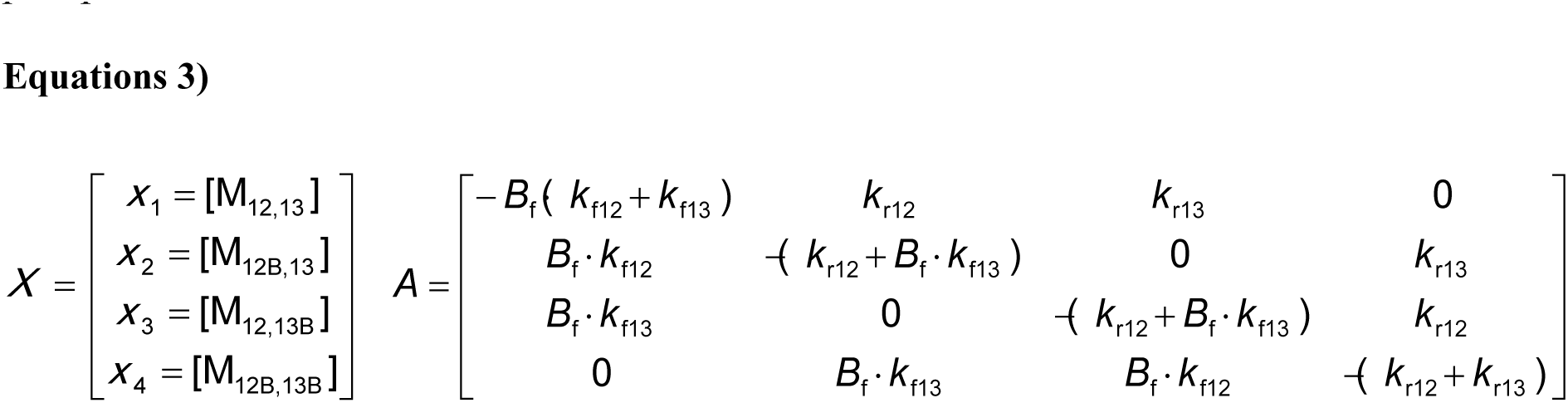

In the above rate matrix *B*_f_ is the concentration of free Bub, and we assume [Bub]_T_ = 30 nM. The equilibrium concentration of each state was obtained by numerically solving 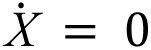.

Following the experimental findings (Figures 4 and 5), we assume that the total concentration of the eSAC activator complex in the cell exceeds the total concentration of Bub. This assumption makes the abundance of Bub-bound eSAC activator complexes dependent on the total concentration of the eSAC activator complex in the cell. At very low concentrations of the eSAC kinase domain, the concentration of eSAC activator complex is much smaller than the total concentration of Bub. Therefore, the eSAC activator complexes tend to be fully loaded, with a Bub on every MELT motif. For example, for a phosphodomain with the MELT motifs, the most abundant complex will be M_12B,13B_ (Figure 5 and Figure S5). However, at high concentration of the eSAC kinase domain, the eSAC activator complexes far outnumber Bub. Consequently, the most abundant eSAC activator complexes either bind a single Bub at one of the MELT motifs, or do not bind any Bub at all. This effect is also evident in the Bub-bindings curves (Figure 5 and Figure S5). It should be noted that as the concentration of the eSAC kinase domain increases to its highest values, the total concentration of phosphorylated MELT motifs bound to Bub increases until it reaches the saturation point equivalent to the total concentration of Bub in the cell or [Bub]_T_. The concentration of all phosphodomains bound by one or more Bub units is defined as the total concentration of the eSAC.
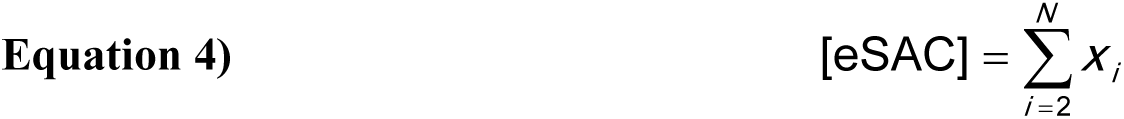

#### Catalysis of Closed-Mad2 conversion by the eSAC phosphodomains

We assume that the recruitment of SAC proteins (Bub in our model) enables the conversion of open-Mad2 into closed/active-Mad2. The rate of this conversion, *k*_amad_, is the concentration-weighted sum of the conversion rates, *k_i_,* of each eSAC activator complex bound by Bub (the suffix *i* denotes the Bub-bound state of the eSAC activator complex):
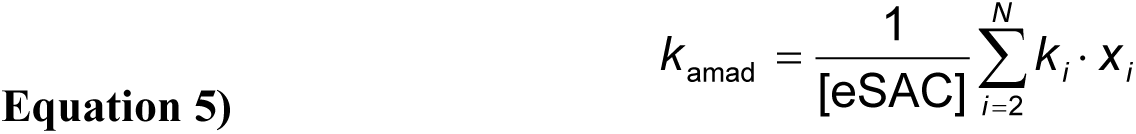

In the above equation, the summation starts from *i* = 2 to exclude the phosphodomains that do not bind Bub. The value of *k_i_* depends on the number of Bub molecules that an eSAC activator complex binds. For phosphodomains containing up to 4 MELT motifs, when the eSAC activator complex binds more than one Bub, the cumulative rate of closed-Mad2 generation was assumed to be additive. Thus, *k*_12B,13B_ = *k*_12B_ + *k*_13B_ for the eSAC phosphodomain that binds a Bub at the 12^th^ and 13^th^ MELT motif Table S2). As discussed above, eSAC activator complexes bound by multiple Bub molecules are the most abundant species at very low [Mps1]. Consequently, *k*_amad_ at very low [Mps1] is dominated by these species. Hence, in this region *k*_amad_ increases with the number of MELT motifs on each phosphodomain. With increasing [Mps1], all the curves decrease monotonically towards their corresponding saturation value, which is essentially determined by the *k*_amad_ value and concentration of states with only one bound Bub. The dependent of the *k*_amad_ rates on the concentration of the various eSAC phosphodomains are plotted in Figure S6B.

As discussed in the manuscript, the simple scheme discussed so far did not accurately predict the dose-response curve observed for the eSAC phosphodomain containing 6 MELT motifs (compare Figure 5B with the dashed black curve in Figure 6D). Therefore, it was necessary to assume that the eSAC activator complex molecules that recruited more than one Bub produced active/closed Mad2 at a rate that is higher than the rate predicted by adding the activities of the individual MELT motifs bound by Bub. These rates are highlighted in red in Table S2. The increase in the *k*_amad_ rate due to synergistic activity is modest: the values used in the model range from 5 to 20% over the baseline rate (Table S2). It should be noted that the choices of Bub binding states for which we assumed cooperativity, and its level (multiplicative factors), are not unique. Different combinations of cooperative states and cooperativity levels will likely result in a similar looking dose-response curve for the eSAC phosphodomain with 6 MELT motifs. However, we did not undertake an exhaustive analysis of the parameter space to find the entire range of permissible parameter values. This is because our aim was to demonstrate that some level of cooperativity between MELT motifs is necessary to explain the experimental data.

#### The effect of eSAC on exit from mitosis

To study the effect of the closed/active Mad2 generated by the eSAC on mitotic exit, we used a model of mitotic checkpoint proposed by (He et al., 2011). The molecular schematic of this model is displayed in Figure S6A. In this model, Cyclin B (symbolized as CycB) is synthesized at a constant rate and degraded by APC-Cdc20-dependent ubiquitination (upper left corner of Figure S6A). The abundance of Cyclin B determines the activity of CDK-CycB complexes. In the original model, the anaphase-inhibitory signal was generated by chromosomes not under tension. We replaced this variable with the eSAC, and assumed, as in the original model, that the strength of the eSAC signal depends positively on CDK-CycB activity and negatively on an unspecified (“counter-acting”) protein phosphatase (CAPP) (Bouchoux and Uhlmann, 2011; Sullivan et al., 2004). This is a reasonable assumption, because the eSAC uses the functional domains of the same two proteins that normally activate the SAC from unattached/tensionless kinetochores.

We adapted this model by replacing unattached kinetochores with the eSAC. An active eSAC generates closed Mad2 as described in the previous section. Closed/active Mad2 binds reversibly to Cdc20, and sequesters it in the form of the MCC. The MCC dissociates, if the bound Mad2 molecule undergoes spontaneous inactivation. We also assume that active APC-Cdc20 promotes the inactivation of Mad2 in mitotic checkpoint complexes. The assumed positive feedback of active Cdc20 on its own release from the MCC provides a means to accelerate the activation of APC:Cdc20 during the transition into anaphase.

The dynamics of our eSAC model are described by the ordinary differential equations:
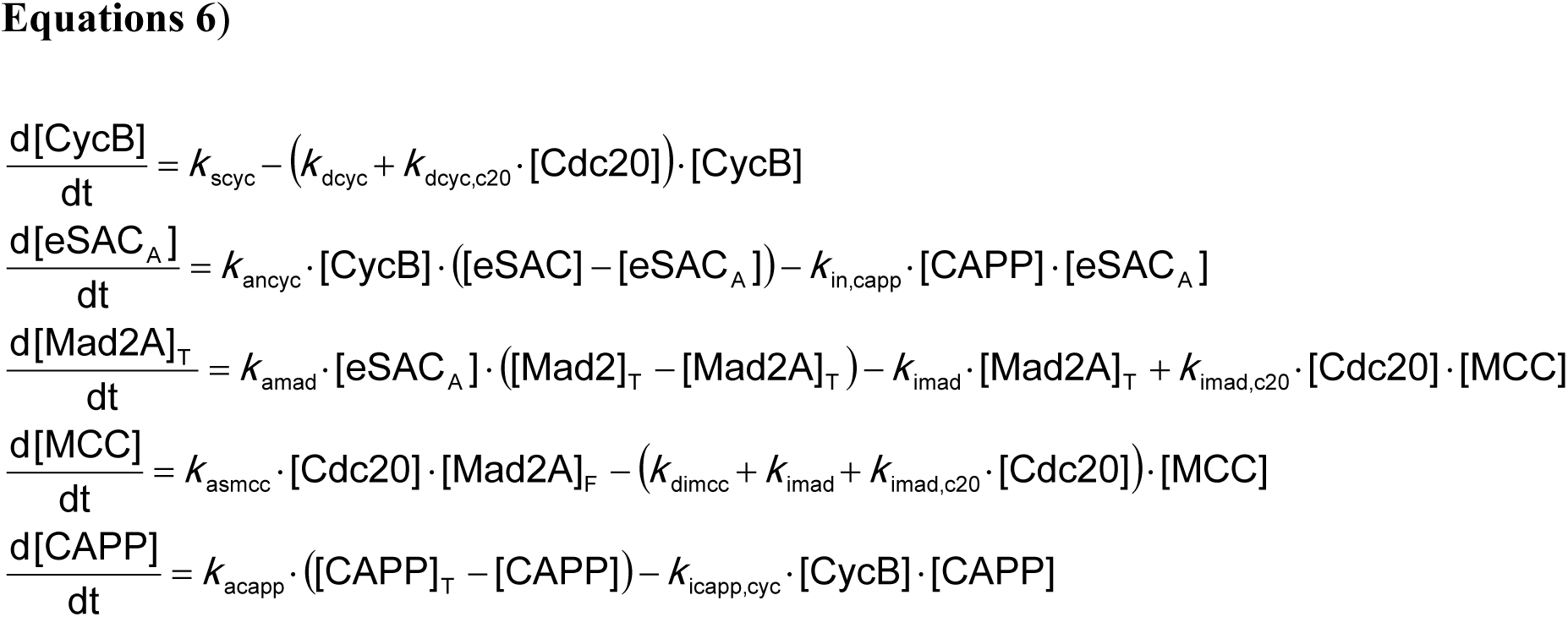

In the above equations, [Name] = concentration of the named species, in arbitrary units. CycB represents the active cyclin-dependent kinase (CDK-CycB) and Cdc20 represents the active anaphase promoting complex (APC-Cdc20). eSAC refers to the dimer between phosphodomain and Mps1 kinase, and we assume that it can be in an actively signaling state eSAC_A_ or an inactive state eSAC_I_ ([eSAC] = [eSAC_A_] + [eSAC_I_]). MCC and CAPP refer to the mitotic checkpoint complex and the CDK counteracting protein phosphatase, respectively. In addition, the total concentration of Mad2 and Cdc20 proteins are:
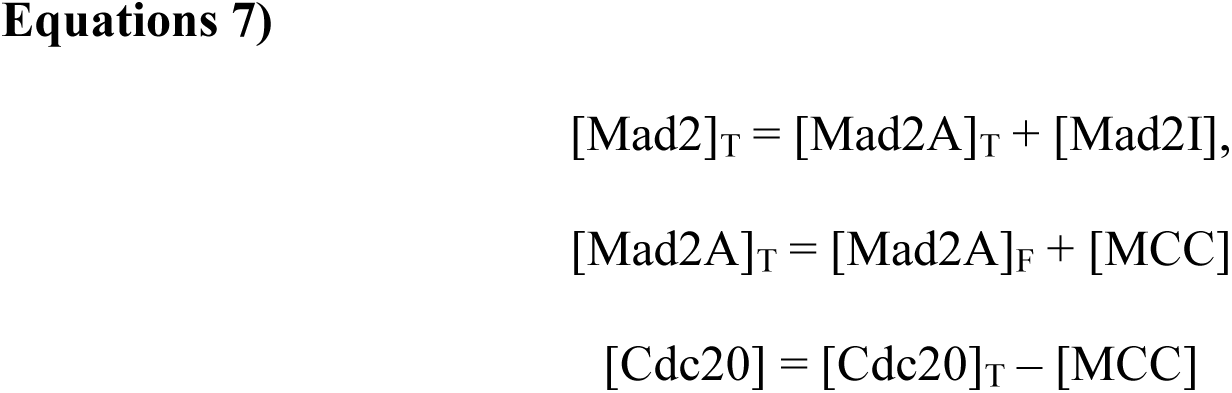

Here, the subscript T indicates the total concentration, subscripts A and I indicate active/closed and inactive/open forms of Mad2, and the subscript F indicates free Mad2A molecules. In the above equations the parameters [eSAC] and *k*_amad_ depend on the Mps1 concentration, as described in the previous section. The values of the other parameters in the model and of the fixed concentrations of some components are listed in Table S1 (He et al., 2011)).

This model of the spindle assembly checkpoint behaves like a bistable switch, which may be either ON (checkpoint engaged, cell blocked in metaphase) or OFF (checkpoint disengaged, cell progresses into anaphase). To illustrate the operating point of this switch, the *k*_amad_ curves for eSAC phosphodomains containing two and six MELT motifs, along with their corresponding bifurcation curves (the values of *k*_amad_ above/below which the switch is ON/OFF) are plotted in Figure S6C. For both phosphodomains, the bifurcation curves are higher than the *k*_amad_ curves, meaning that the switch is OFF in both cases. Hence, in both cases the cell will exit mitosis only after the Cyclin B level in the cell drops below a threshold value. Because the difference between the actual *k*_amad_ value and the bifurcation point is small, the checkpoint control system starts close to bifurcation point where the rates of change of all variables are very small.

#### Simulation of time in mitosis

We explicitly simulated the ‘time in mitosis’ as follows. We assumed that cells remain in mitosis as long as the Cyclin B concentration in the cell is > 5 a.u. (arbitrary units). Therefore, the time at which the concentration falls below this threshold was defined as the time in mitosis. We numerically integrated the ODEs described in the previous section to calculate the time evolution of different species. Typical time courses for CycB concentration for different eSAC phosphodomains, for [eSAC] = 10 a.u., are displayed in Figure S6D. Since the operating point of the bistable switch is OFF, the system always comes out of mitosis (as seen by the drop in [CycB]), albeit after different time delays.

The time evolution of [CycB] can be understood as follows. Initially, CycB is degraded at a very slow rate because the concentration of free Cdc20 is low and the rate of Cdc20-APC mediated degradation of CycB is only slightly larger than the rate of CycB synthesis. The slow rate of CycB degradation can also be understood from our observation that the system starts off close to the bifurcation point, where the rates of change of all quantities are small (Figure S6C). As [CycB] falls below the assumed threshold value, two positive feedback loops get activated, causing a catastrophic drop in [CycB]. The first feedback loop negatively impacts the generation of MCC by inactivating the eSAC, likely by preventing it from recruiting SAC proteins. A lower rate of MCC generation means an increase in [Cdc20], and hence increased degradation of Cyclin B. The second feedback loop impacts the disassembly of MCC by active APC:Cdc20 complexes.

It is important to note here that the system operates close to the bifurcation point (Figure S6C). This operating point suggests that its response (measured by the time in mitosis) will be noisy, as small changes in parameter values can result in a large change in response. This prediction is indeed consistent with experimental data where significant noise is observed in the time spent by cells in mitosis for all types of phosphodomains (see the dose-response data in Figures 4 and 5). To study the effects of noise on time spent in mitosis will require a reasonable model of stochastic fluctuations all along the pathway, from activation of eSAC domains to the eventual activation of APC:Cdc20 complexes. At this stage, we have restricted our efforts to analyzing the average behavior of ‘time in mitosis’ using deterministic simulations.

---

**Figure S1 (Related to Figure 1).**
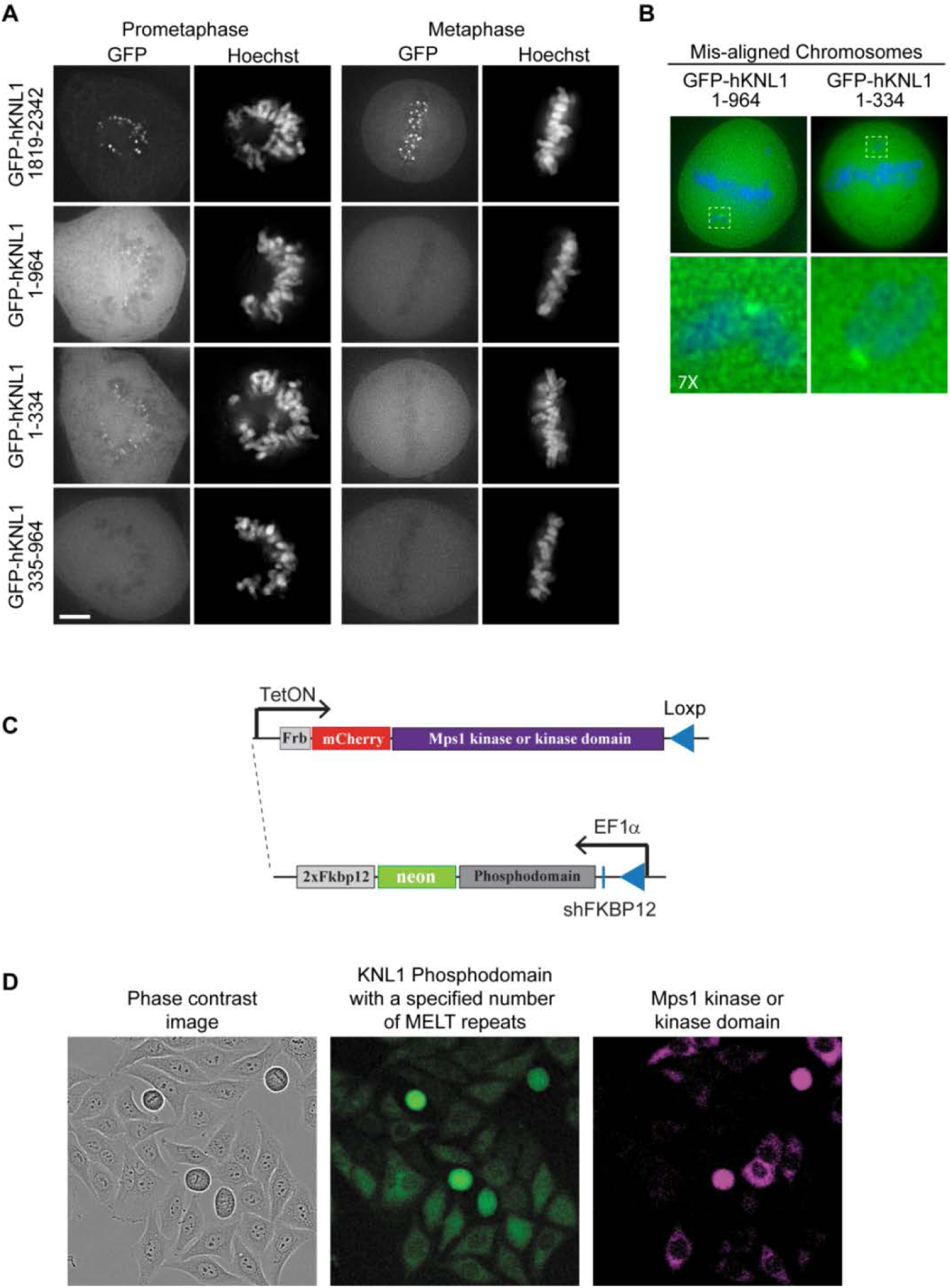
The design and implementation of the ectopic SAC activator. (A) Fluorescent images showing the localization of the indicated KNL1 regions as GFP fusions in live HeLa cells (numbers alongside each micrograph denote the residues in each construct). As expected, the known kinetochore-binding domain at the C-terminus of KNL1 strongly localized to kinetochores. Surprisingly, residues proximal to its N-terminus also transiently localized to kinetochores only in prometaphase (Scale bar = 5 μm). (B) Images at the top display metaphase cells with misaligned chromosomes. KNL1 fragments containing it N-terminus also localized to kinetochores on these chromosomes. Dashed boxes highlight the magnified areas shown in the bottom right panel. (C) Each eSAC cell line was created by integrating a bi-cistronic cassette in the HeLa genome at a unique Loxp site via Cre-mediated recombination. This strategy was developed in Ballister et al 2013. (B) The Phosphodomain-neonGFP-2xFkbp12, wherein the phosphodomain cassette can contain a specified numbers of MELT motifs, is constitutively expressed by the EF1α promoter. The 5’ UTR of this gene also contains a sequence encoding shRNA against the endogenous FKBP protein. Frb-mCherry-Mps1, wherein Mps1 can be either the full length gene or just the sequence encoding its kinase domain is expressed conditionally from a TetON promoter.

**Figure S2 (Related to Figure 2).**
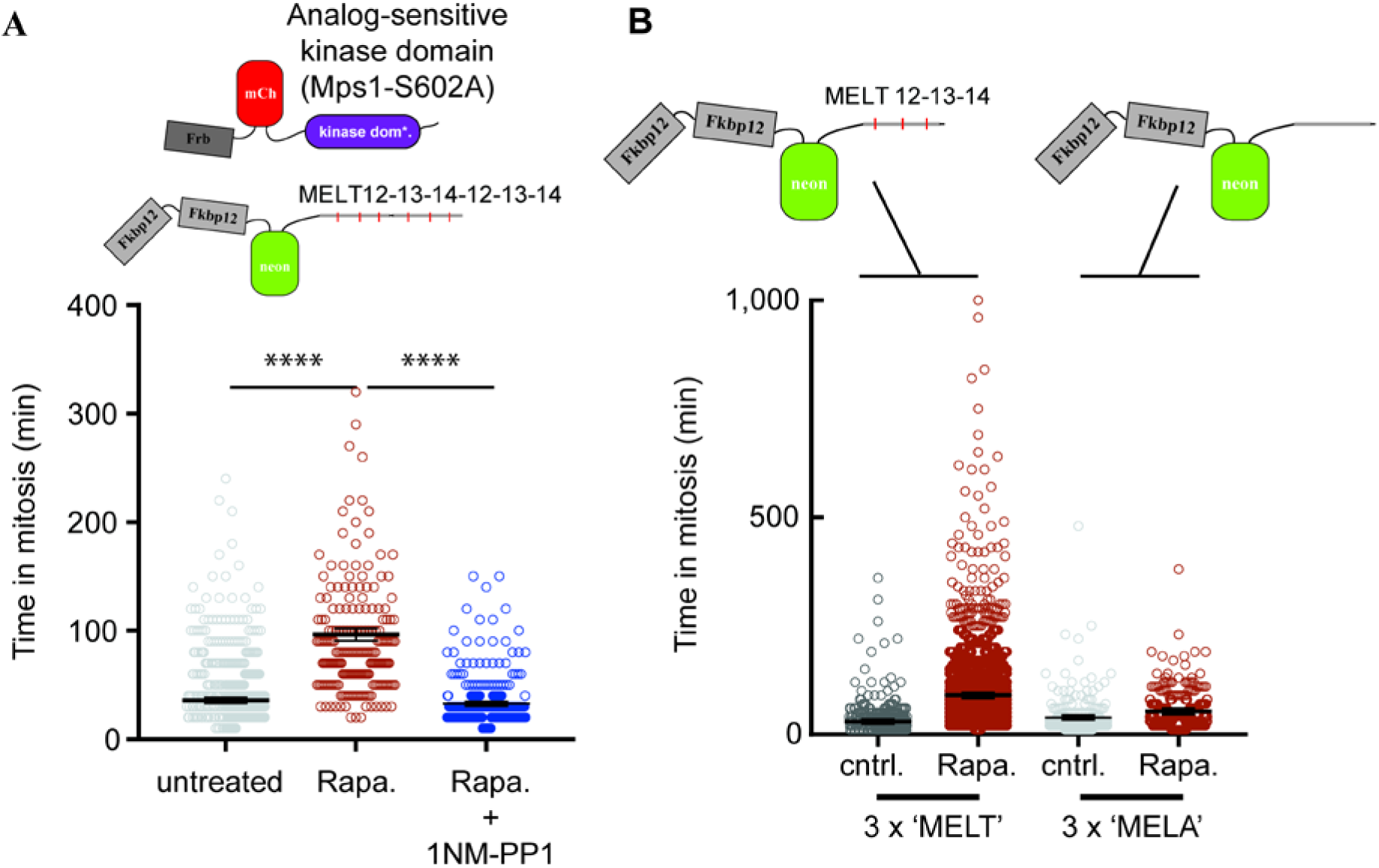
Phosphorylation of the MELT motifs in the minimal KNL1 phosphodomain by the Mps1 kinase domain is necessary for eSAC activity. (A) Kinase activity of the Mps1 kinase domain is necessary for the rapamycin-induced mitotic arrest. Rapamycin-induced dimerization of the analog-sensitive allele of the Mps1 kinase domain, Mps1S602A, with the minimal phosphodomain (displayed in the cartoon at the top) produced a weak mitotic arrest. The weak eSAC-induced arrest likely indicates a significantly reduced activity of the mutant kinase domain. Combined treatment with Rapamycin and the ATP analog 1NM-PP1 (10 μM) abrogated the eSAC-induced arrest. Cells expressing > 5 a. u. of Frb-mCherry-Mps1S602A were used for this analysis (n=177, 206, and 332 respectively; p<0.0001, Mann-Whitney test). (B) Phosphorylatable MELT motifs are necessary for rapamycin-induced mitotic arrest. The minimal phosphodomain used in the experiment is displayed at the top. n = 853, 2238, 252, and 313 respectively. In both A and B, black horizontal lines display mean ± s.e.m.

**Figure S3 (Related to Figures 4 and 5).**
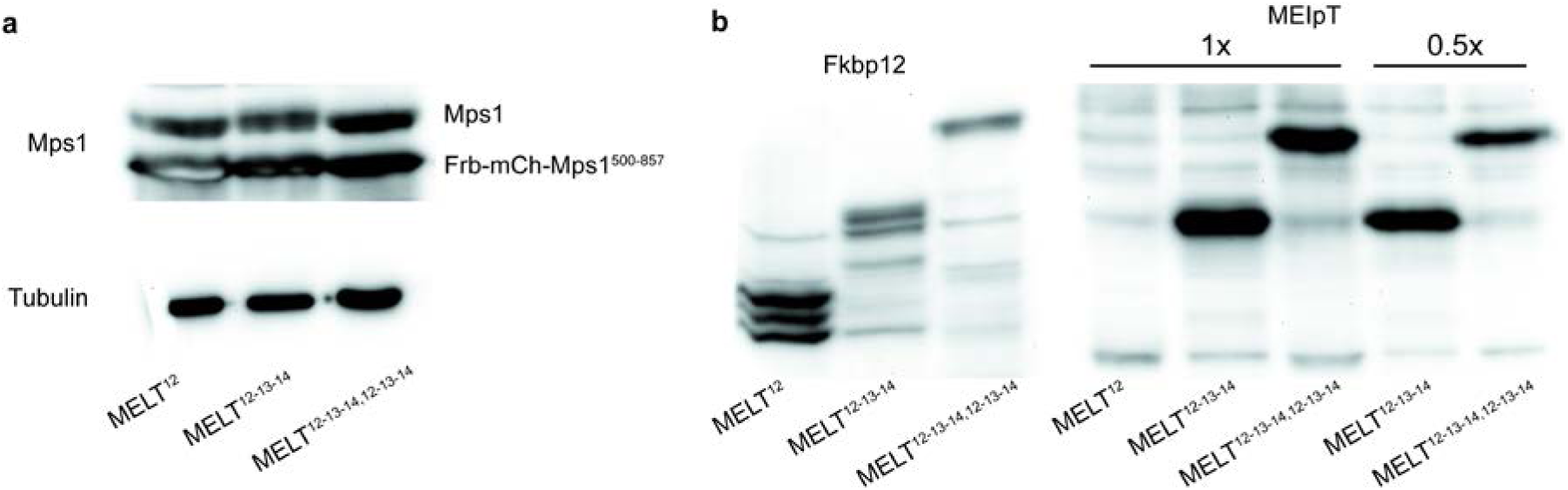
The abundance of the eSAC kinase domain is comparable with that of the endogenous Mps1 kinase. (**A**) Comparison of the expression level of the endogenous Mps1 and the eSAC Mps1 kinase domain (Frb-mCherry-Mps1^500-857^) for three different cell lines expressing the indicated phosphodomains. Total cell lysates probed with an antibody against the C-terminus of Mps1, which is present in the kinase domain used for the eSAC. (**B**) Assessment of the expression levels of the eSAC phosphodomains (antibody against the Fkbp12 protein) and their phosphorylation after rapamycin-induced dimerization with Mps1 (phosphospecific antibody against the 13^th^ MELT motif does not recognize the phosphodomain containing the 12^th^ MELT motif alone).

**Figure S4 (Related to Figures 4 and 5).**
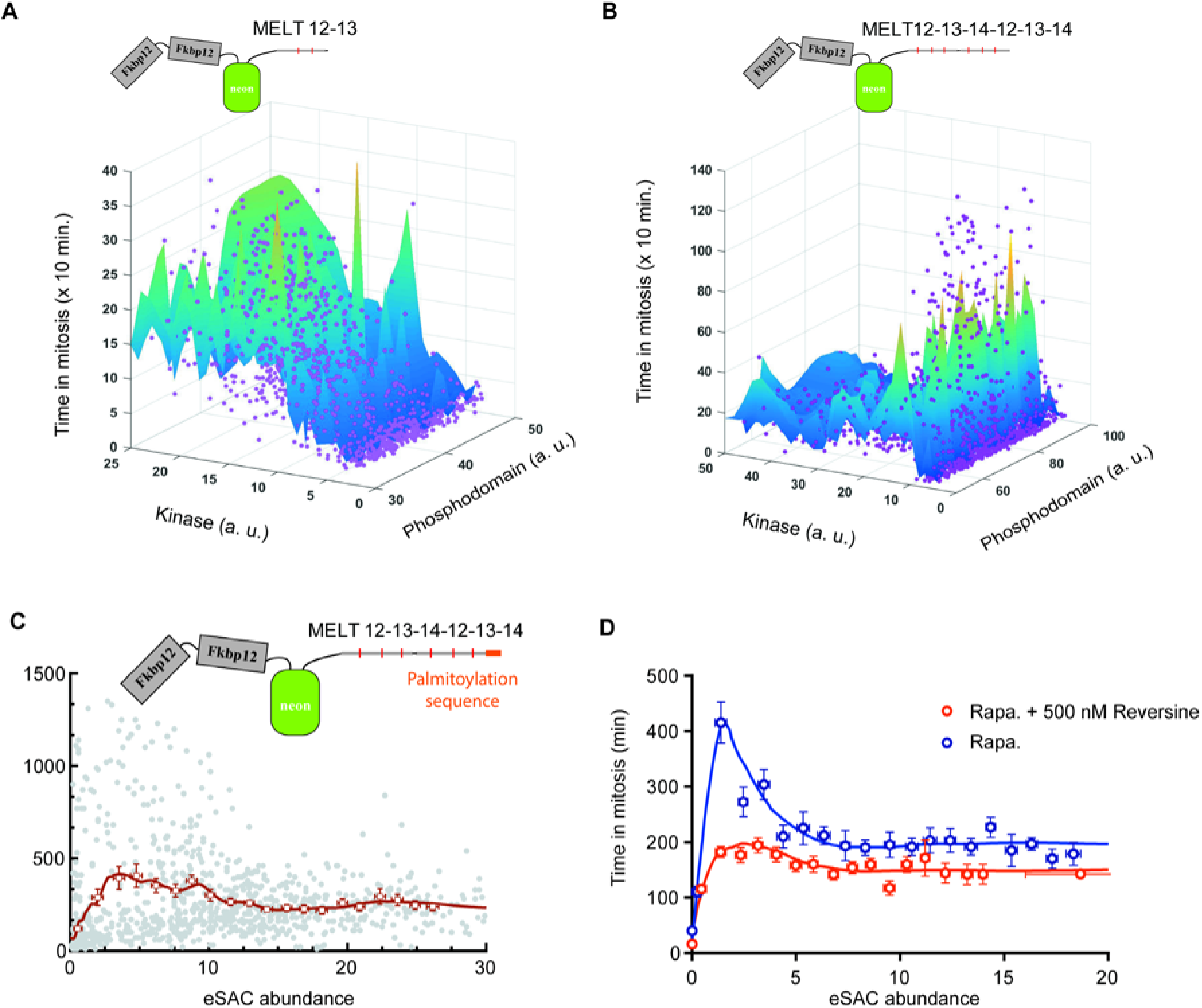
The duration of the eSAC-induced metaphase arrest correlates strongly with the cellular abundance of the kinase domain, but not with the abundance of the phosphodomain. (A-B) Dependence of eSAC-induced mitotic duration on the abundance of the phosphodomain (neonGreen fluorescence) and Mps1 kinase domain (mCherry fluorescence) shown for minimal KNL1 phosphodomains containing 2 and 6 MELT motifs respectively (n = 1376 for a and n = 1877 for b). In A and B, magenta spots represent measurements from one cell. The surface was calculated using the ‘griddata’ function with cubic interpolation in MatLab. (C) Dose-response characteristics of a phosphodomain containing 6 MELT motifs when it is targeted to the plasma membrane (n=1056). (D) The complex nature of the dose-response characteristics is retained even when the endogenous SAC activation switch is inactivated by inhibiting the kinase activity of the endogenous Mps1. Open squares and circles represent averages of binned data, error bars represent s.e.m.

**Figure S5 (Related to Figure 6).**
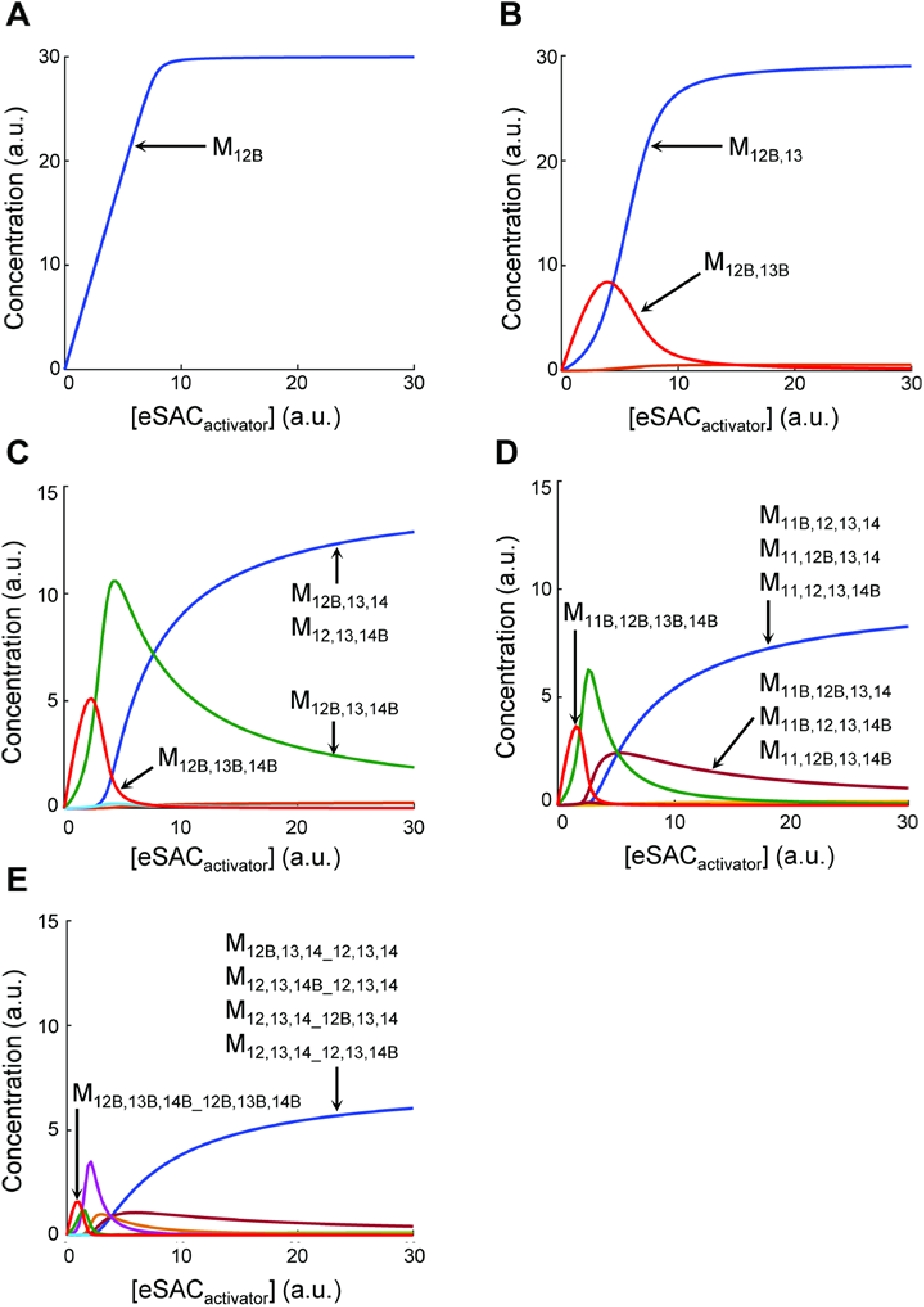
Changes in the abundance of Bub-bound species at different concentrations of the eSAC activator complexes. (A-E) The abbreviations for different Bub-bound phosphodomain species are as follows. The subscripted number following the M denotes the rank of the MELT motif in the KNL1 phosphodomain (see Fig. 4A). For example, M_12_ symbolizes the eSAC phosphodomain with one MELT motif, and M_12,13,14_12,13,14_ symbolizes the phosphodomain with six MELT motifs. The subscript ‘B’ in front of the number signifies that the MELT motif denoted by the number is bound by Bub. The concentration of Bub is assumed to be 30 a.u.

**Figure S6 (Related to Figure 6).**
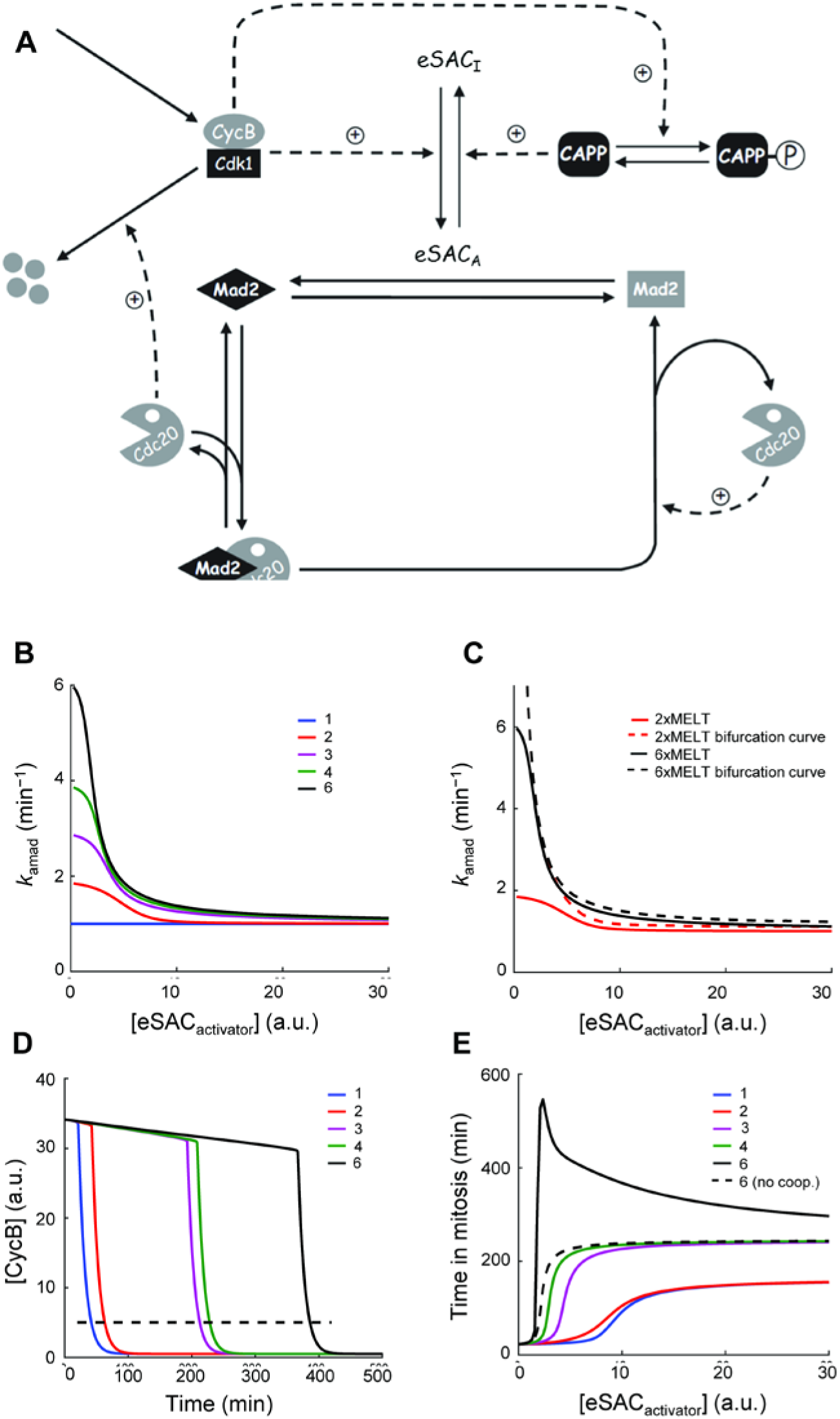
Schematic of the model used to simulate anaphase onset (adapted from ref. (He et al., 2011)). **(A)** An active eSAC produces the active or “closed” form of Mad2, which sequesters Cdc20 as part of the MCC. This eSAC activity is promoted by high CyclinB-CDK1, and inhibited by an unspecified phosphatase (CAPP). High CyclinB-CDK1 activity also inhibits the activity of the phosphatase. eSAC catalyzes the formation of the active/closed Mad2 at a rate *k_amad_* determined by: (1) the amino acid sequences of the MELT motifs in the eSAC phosphodomain, (2) synergistic activity of the MELT motifs, and (3) abundance of the eSAC activator. Closed/active Mad2 sequesters Cdc20, and thus inhibits the APC. Free Cdc20 acts with APC to degrade Cyclin B and to promote the dissociation of the Mad2-Cdc20 complex. Thus, *k_amad_* also determines the rate of degradation of Cyclin B by the APC. When Cyclin B levels fall below the minimum threshold value, the two feedback loops work concurrently to rapidly drive the cell out of mitosis. (**B**) Dependence of the cumulative rate of generation of active/closed Mad2, *k_amad_,* by different eSAC phosphodomain plotted on the eSAC abundance. (**C**) *k_amad_* curves for eSAC phosphodomains containing 2 and 6 MELT motifs, along with their corresponding bifurcation curves (*k_amad_* values above/below which the switch is ON/OFF). (**D**) Change in Cyclin B concentration with respect to time for different eSAC phosphodomains. We assume that the cell exits mitosis when the Cyclin B concentration falls below 5 a.u. (dashed line). (**E**) Dependence of time in mitosis as a function of the eSAC activator complex concentration for different eSAC phosphodomains. The dashed curve corresponds to dose response curve for the phosphodomain containing 6 MELT motifs calculated by assuming that cooperativity is absent.

**Table S1 (related to Figure 6).** Parameter values used in the simulation of the eSAC

| Parameter | Value | Parameter | Value |
| --- | --- | --- | --- |
| $k_{f11}$ | $1 \text{ nM}^{-1} \text{ min}^{-1}$ | $k_{r11}$ | $0.1 \text{ min}^{-1}$ |
| $k_{f12}$ | $1 \text{ nM}^{-1} \text{ min}^{-1}$ | $k_{r12}$ | $0.1 \text{ min}^{-1}$ |
| $k_{f13}$ | $1 \text{ nM}^{-1} \text{ min}^{-1}$ | $k_{r13}$ | $5 \text{ min}^{-1}$ |
| $k_{f14}$ | $1 \text{ nM}^{-1} \text{ min}^{-1}$ | $k_{r14}$ | $0.1 \text{ min}^{-1}$ |
| $k_{scys}$ | $0.001 * [\text{CycB}]_M \text{ min}^{-1}$ | $k_{dimcc}$ | $1 \text{ min}^{-1}$ |
| $k_{dcyc}$ | $0.001 \text{ min}^{-1}$ | $k_{acapp}$ | $0.1 \text{ min}^{-1}$ |
| $k_{dcyc,c20}$ | $0.1 / [\text{Cdc20}]_T \text{ min}^{-1}$ | $k_{icapp,cyc}$ | $1 / [\text{CycB}]_M \text{ min}^{-1}$ |
| $k_{an,cyc}$ | $1 \text{ nM}^{-1} \text{ min}^{-1}$ | $[\text{CycB}]_M$ | 50 nM |
| $k_{in,capp}$ | $1 \text{ nM}^{-1} \text{ min}^{-1}$ | $[\text{Cdc20}]_T$ | 25 nM |
| $k_{imad}$ | $0.1 \text{ min}^{-1}$ | $[\text{Mad2}]_T$ | 50 nM |
| $k_{imad,c20}$ | $10 / [\text{Cdc20}]_T \text{ min}^{-1}$ | $[\text{CAPP}]_T$ | 50 nM |
| $k_{asmcc}$ | $400 / [\text{Cdc20}]_T \text{ min}^{-1}$ | | |

**Table S2 (Related to Figure 6).** Rates of open-Mad2 to closed/active-Mad2 conversion by the Bub binding states. All the rates were multiplied by a scaling factor 0.013 in order to scale the simulation results to match the experimental data. For the sake of brevity, we have shortened the MELT motif numbering notation using the rule: 11->1, 12->2, 13->3, 14->4. Only the numbers of Bub-bound MELT motifs are listed. In the case of eSAC phosphodomain with 6 MELT motifs the underscore symbol is used to differentiate between the two sets of MELT 12,13,14. Cooperative interactions are highlighted in red.

| Parameter | Value | Parameter | Value | Parameter | Value |
| --- | --- | --- | --- | --- | --- |
| $k_4$ | 1.02 | $k_{2\_2}$ | $2 * k_2$ | $k_{34\_34}$ | $2 * k_{34}$ |
| $k_3$ | 1 | $k_{234}$ | $k_2 + k_3 + k_4$ | $k_{34\_24}$ | $k_{34} + k_{24}$ |
| $k_2$ | 1 | $k_{4\_34}$ | $k_4 + k_{34}$ | $k_{34\_23}$ | $k_{34} + k_{23}$ |
| $k_1$ | 1.01 | $k_{4\_24}$ | $k_4 + k_{24}$ | $k_{2\_234}$ | $k_2 + k_{234}$ |
| $k_{34}$ | $k_3 + k_4$ | $k_{4\_23}$ | $k_4 + k_{23}$ | $k_{24\_34}$ | $k_{24} + k_{34}$ |
| $k_{24}$ | $k_2 + k_4$ | $k_{3\_34}$ | $k_3 + k_{34}$ | $k_{24\_24}$ | $2 * k_{24} * 1.08$ |
| $k_{23}$ | $k_2 + k_3$ | $k_{3\_24}$ | $k_3 + k_{24}$ | $k_{24\_23}$ | $k_{24} + k_{23}$ |
| $k_{14}$ | $k_1 + k_4$ | $k_{3\_23}$ | $k_3 + k_{23}$ | $k_{23\_34}$ | $k_{23} + k_{34}$ |
| $k_{13}$ | $k_1 + k_3$ | $k_{34\_4}$ | $k_{34} + k_4$ | $k_{23\_24}$ | $k_{23} + k_{24}$ |
| $k_{12}$ | $k_1 + k_2$ | $k_{34\_3}$ | $k_{34} + k_3$ | $k_{23\_23}$ | $2 * k_{23}$ |
| $k_{123}$ | $k_1 + k_2 + k_3$ | $k_{34\_2}$ | $k_{34} + k_2$ | $k_{234\_4}$ | $k_{234} + k_4$ |
| $k_{124}$ | $k_1 + k_2 + k_4$ | $k_{2\_34}$ | $k_2 + k_{34}$ | $k_{234\_3}$ | $k_{234} + k_3$ |
| $k_{134}$ | $k_1+k_3+k_4$ | $k_{2\_24}$ | $k_2+k_{24}$ | $k_{234\_2}$ | $k_{234}+k_2$ |
| $k_{1234}$ | $k_1+k_2+k_3+k_4$ | $k_{2\_23}$ | $k_2+k_{23}$ | $k_{34\_234}$ | $k_{34}+k_{234}$ |
| $k_{4\_4}$ | $2*k_4*1.2$ | $k_{24\_4}$ | $k_{24}+k_4$ | $k_{24\_234}$ | $(k_{24}+k_{234})*1.05$ |
| $k_{4\_3}$ | $k_4+k_3$ | $k_{24\_3}$ | $k_{24}+k_3$ | $k_{23\_234}$ | $k_{23}+k_{234}$ |
| $k_{4\_2}$ | $k_4+k_2$ | $k_{24\_2}$ | $k_{24}+k_2$ | $k_{234\_34}$ | $k_{234}+k_{34}$ |
| $k_{3\_4}$ | $k_3+k_4$ | $k_{23\_4}$ | $k_{23}+k_4$ | $k_{234\_24}$ | $(k_{234}+k_{24})*1.05$ |
| $k_{3\_3}$ | $2*k_3$ | $k_{23\_3}$ | $k_{23}+k_3$ | $k_{234\_23}$ | $k_{234}+k_{23}$ |
| $k_{3\_2}$ | $k_3+k_2$ | $k_{23\_2}$ | $k_{23}+k_2$ | $k_{234\_234}$ | $2*k_{234}*1.05$ |
| $k_{2\_4}$ | $k_2+k_4$ | $k_{4\_234}$ | $k_4+k_{234}$ | | |
| $k_{2\_3}$ | $k_2+k_3$ | $k_{3\_234}$ | $k_3+k_{234}$ | | |

**Supplemental Movie S1**

Representative movie displays time-lapse imaging of DMSO-treated eSAC cell line. DNA (magenta) is labeled with low concentration of Hoechst and the microtubules (green) are labeled with siR-Tubulin. Numbers on the lower right corner indicate elapsed time [hr:min].

**Supplemental Movie S2**

Representative movie displays time-lapse imaging of rapamycin-treated eSAC cell line. DNA (magenta) is labeled with low concentration of Hoechst and the microtubules (green) are labeled with siR-Tubulin. Numbers on the lower right corner indicate elapsed time [hr:min].

**Supplemental Movie S3**

Representative movie displays long-term time-lapse image of an untreated eSAC cells expressing eSAC phosphodomain with 3 MELT motifs. Each frame is an overlay of the phase contrast and mCherry (reporter for the eSAC kinase domain). Numbers in the lower left hand corner display time [hr:min].

**Supplemental Movie S4**

Representative movie displays long-term time-lapse image of rapamycin eSAC cells expressing eSAC phosphodomain with 4 MELT motifs. Each frame is an overlay of the phase contrast and mCherry (reporter for the eSAC kinase domain). Numbers in the lower left hand corner display time [hr:min].

**Supplemental Movie S5**

Representative movie displays long-term time-lapse image of rapamycin eSAC cells expressing eSAC phosphodomain with 1 MELT motif. Each frame is an overlay of the phase contrast and mCherry (reporter for the eSAC kinase domain). Numbers in the lower left hand corner display time [hr:min].

**Supplemental Movie S6**

Representative movie displays long-term time-lapse image of rapamycin eSAC cells expressing eSAC phosphodomain with 6 MELT motifs. Each frame is an overlay of the phase contrast and mCherry (reporter for the eSAC kinase domain). Numbers in the lower left hand corner display time [hr:min].

